# Multi-omic Characterization of Human Tubular Epithelial Cell Response to Serum

**DOI:** 10.1101/2021.01.29.428186

**Authors:** Kevin A. Lidberg, Selvaraj Muthusamy, Mohamed Adil, Ranita S. Patel, Lu Wang, Theo K. Bammler, Jonathan Reichel, Catherine K. Yeung, Jonathan Himmelfarb, Edward J. Kelly, Shreeram Akilesh

## Abstract

Proteinuria, the spillage of serum proteins into the urine, is a feature of glomerulonephritides, podocyte disorders and diabetic nephropathy. However, the response of tubular epithelial cells to serum protein exposure has not been systematically characterized. Using transcriptomic profiling we studied serum-induced changes in primary human tubular epithelial cells cultured in 3D microphysiological devices. Serum proteins induced cellular proliferation, cytokine secretion and activated a coordinated stress response. We orthogonally confirmed our findings by comparing the transcriptomic and epigenomic landscapes of intact human kidney cortex and isolated tubular epithelial cells cultured in fetal bovine serum. Importantly, key transcriptomic programs in response to either type of serum exposure remained consistent, including comparisons to an established mouse model of kidney injury. This serum-induced transcriptional response was dominated by switching off of nuclear receptor-driven programs and activation of AP-1 and NF-κB signatures in the tubular epigenomic landscape. These features of active regulation were seen at canonical kidney injury genes (*HAVCR1*) and genes associated with COVID-19 (*ACE2*, *IL6*). Our data provide a reference map for dissecting the regulatory and transcriptional response of kidney tubular epithelial cells injury induced by serum.

## Introduction

Chronic kidney disease (CKD) describes a broad range of conditions characterized by a sometimes progressive and irreversible loss of renal function that can result in end-stage kidney disease (ESKD). CKD and ESKD are a major health concern because they decrease quality of life, increase morbidity and mortality, and place a considerable economic burden on the US healthcare system (*1–3*). While diabetes and hypertension are the primary risk factors for developing CKD, episodes of acute kidney injury (AKI), especially if severe or recurrent, can increase risk and worsen disease (*4*). Vascular rarefaction, tubular atrophy, interstitial inflammation and fibrosis are the histological hallmarks associated with loss of kidney function in CKD. However, these histological changes occur regardless of the underlying cause of CKD suggesting that processes independent of the initiating disorder can drive disease progression.

Proteinuria, the spillage of serum proteins into the urine due to a loss of glomerular barrier selectivity, occurs in most forms of CKD and is associated with disease pathogenesis. Proteinuria is a prognostic biomarker for CKD progression and its reduction is associated with favorable clinical outcomes, particularly when baseline proteinuria is highest (*5–7*). In both humans with glomerular disease (e.g., lupus glomerulonephritis) and animals with experimentally derived glomerular dysfunction, the degree of proteinuria correlates with the severity of tubular lesions (*8–10*). Rather than being correlative, the association between proteinuria and pathophysiological tubular changes may be causative, as several lines of evidence support a role for a direct action of proteinuria on the proximal tubule. First, proximal tubule epithelial cells (PTECs) induce and secrete vasoactive and pro-inflammatory factors when treated *in vitro* with serum proteins such as albumin, transferrin, and IgG (*11–13*). Second, the proximal tubules from proteinuric patients exhibit a pro-inflammatory phenotype (*14–16*). Lastly, PTECs reabsorb nearly all protein from the glomerular ultrafiltrate via receptor mediated endocytosis (e.g., megalin-cubilin complex); a function that modulates proteinuria-induced proximal tubule injury in animal models (*17, 18*). As such, proteinuria is thought to play an important role in the formation of tubular lesions, in part through its action on the proximal tubule (*19*).

The proximal tubule is the most active, vascularized, and energy-demanding segment of the nephron responsible for several significant processes (*20*). This aspect makes it a vulnerable target for ischemic, obstructive, drug-induced, and immune-mediated injury. After injury, PTECs mount a repair response wherein the cells dedifferentiate to a proliferation-competent state, replicate, then differentiate to regenerate the epithelium (*21–23*). However, during this process a subset of the PTEC pool exhibits certain attributes such as defective fatty acid metabolism, enhanced pro-inflammatory signaling, and aberrant cell cycle arrest, which are maladaptive if the normal epithelium is not restored (*24–26*). Importantly, injury to the proximal tubule is alone sufficient to cause renal inflammation and fibrosis (*27*). The signaling pathways and epigenomic regulation that control PTEC phenotype during injury or in states of disease (e.g., proteinuria) are therefore of considerable interest as they represent potential therapeutic targets for slowing CKD progression.

Accordingly, we set out to identify the transcriptional programs and epigenetic signatures that control PTEC response to proteinuria using RNA-seq and ATAC-seq and two independent model systems. First, we used a microphysiological system (MPS) where primary human PTECs were treated with serum-free medium or 2% normal human serum. Culturing PTECs in a microfluidic device creates a three dimensional *in vivo*-like microenvironment that enables the cells to better emulate the metabolic, regulatory, and transport activities of the proximal tubule as shown by our group (*28–32*) as well as others (*33–35*). The second approach compared native renal cortex to primary PTECs cultured in 2D in the presence of 10% fetal bovine serum (FBS). This approach allowed us to use the normal *in vivo* regulatory landscape and transcriptome as a reference to identify signatures that are lost due to *in vitro* culture. Finally, we validated key findings from our multi-omic studies by comparing the transcriptomic signature we derived to a widely used mouse model of kidney injury and also by quantifying pro-inflammatory protein secretion in MPS treated with either serum or albumin.

## Results

### Serum exposure induces proliferation in 3D cultures of kidney tubules

Seeding of primary human tubular epithelial cells into the central lumen of a collagen I matrix within a microphysiological device (MPS) reproducibly produces a polarized monolayer of cells with a central lumen (**Figure 1A**). Serum-free growth medium can be continuously perfused through the central channel and cell-conditioned effluent can be collected for measurements. Once established, the tubular epithelial monolayer can be maintained under continuous flow without evidence of tubular epithelial turnover or proliferation as measured by Ki-67 immunofluorescence. In contrast, exposure of cells to 2% serum in the perfusate resulted in an increase in mitotic activity (**Figure 1B, C**).

**Figure 1.**
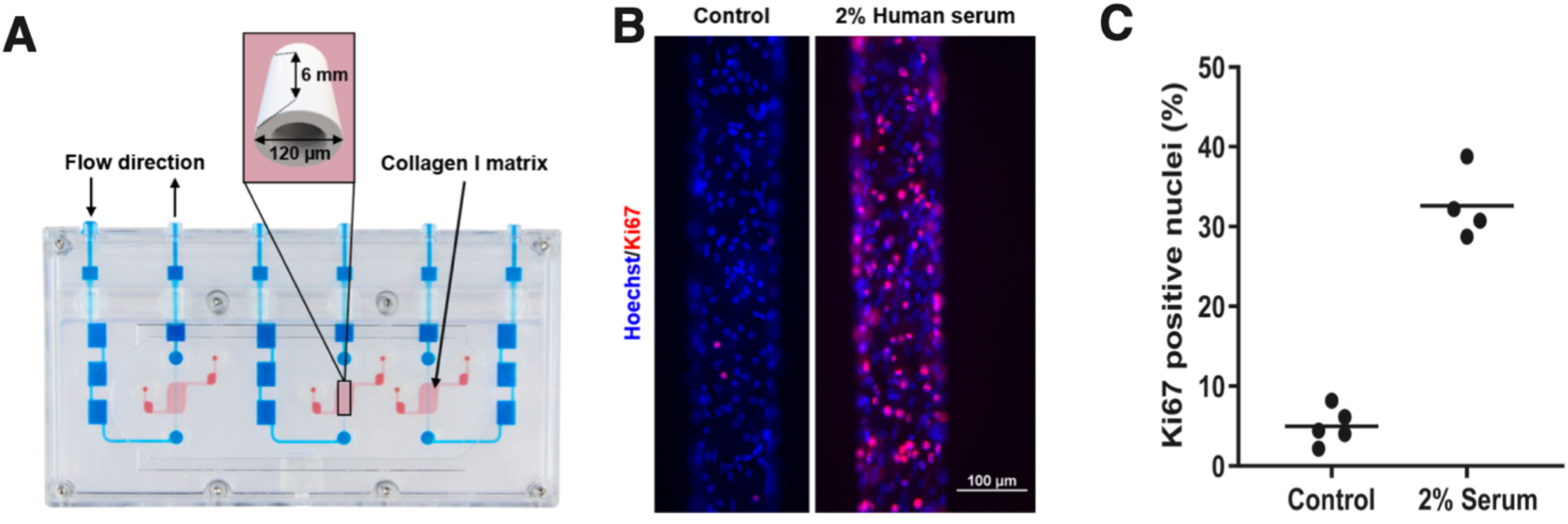
Human kidney tubules in 3D microphysiologic devices proliferate in response to serum exposure. A) Schematic of 2-channel Nortis microphysiologic system (MPS). The red areas represent a 3D matrix chamber through which a central tubule monolayer lined channel can be perfused with medium (blue). B) Compared to control, 2% human serum treated tubule MPS have higher frequency of nuclei that label with the proliferative marker Ki-67. C) Quantification of percent Ki-67 positive nuclei in n=5 control and n=4 serum treated tubular MPS from Donors 1 and 2; p<0.0001 by two-tailed t-test.

### Serum exposure induces cytokine production and metabolic reprogramming in kidney tubule cells

To characterize the changes induced by serum exposure, we extracted RNA from multiple replicates of control or serum-treated tubular MPS, and measured gene expression by RNA-seq. Serum treatment induced upregulation of 533 genes and downregulation of 406 genes with a fold change>1.5 and DESeq2 adjusted p-value<0.05 (**Figure 2A**). Gene ontology enrichment analysis of these differentially expressed genes showed significant upregulation of biological processes related with cytokine-mediated signaling pathways (GO:0019221), extracellular matrix organization (GO:0030198) and negative regulation of apoptotic processes (GO:0043066). Conversely, biological processes associated with regulation of cholesterol metabolic process (GO:0090181) and regulation of alcohol biosynthetic process (GO:1902930) were significantly downregulated (**Figure 2B**). Taken together, exposure of primary human tubular MPS to serum induced expression of genes related to cellular proliferation, extracellular matrix reorganization, inflammatory cytokine secretion with concomitant downregulation of genes related to metabolism and solute transport (**Figure 2C**). Advaita iPathwayGuide analysis identified TNF, EGF, FOXM1 and IL1A/B as key upstream regulators based on differential expression of their known target genes (**Figure 2D**). Pathway analysis identified that cytokine/chemokine-mediated signaling (p=1.120×10^−8^), TNF-(p=4.465×10^−6^) and NF-κB-(p=4.457×10^−6^) mediated signaling pathways were prominent points of regulation in serum-treated tubule MPS (**Figure 2E**).

**Figure 2.**
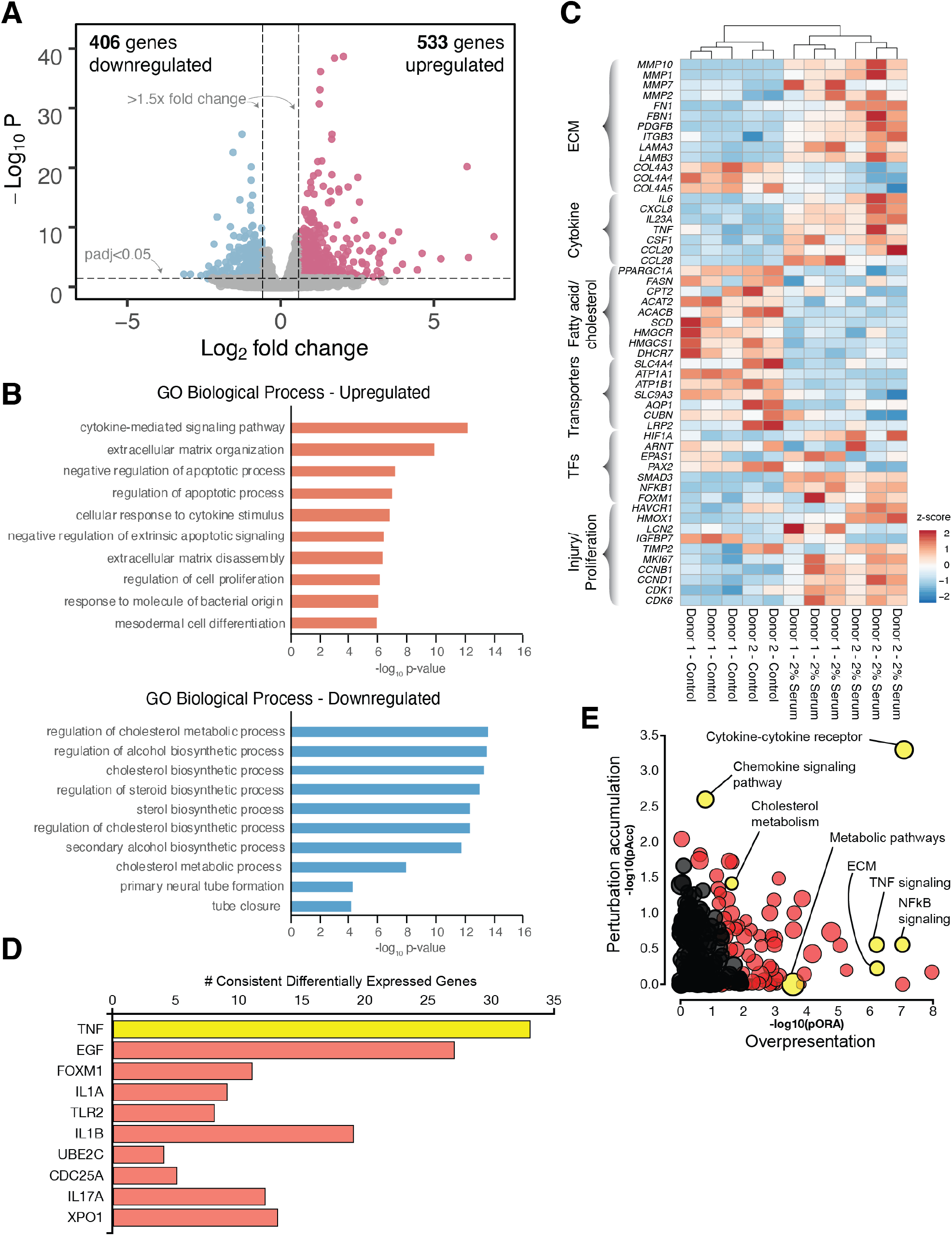
Serum exposure induces cytokine production and metabolic reprogramming in kidney tubule cells. A) Volcano plot of differentially expressed genes in tubule MPS exposed for 48hrs to 2% human serum versus untreated control. Upregulated = higher expression in serum treated MPS. B) Gene ontology enrichments for top 10 up- and down-regulated biological process in the differentially expressed gene set. C) Heatmap of exemplar genes associated with significant gene ontology enrichments. D) Advaita analysis of key upstream regulators showing the number of differentially expressed target genes detected for each indicated upstream regulator. E) Advaita pathway analysis demonstrates significant perturbation of cytokine, chemokine, TNF and NF-κB signaling pathway components in serum-exposed tubule cells. Dots representing pathways are positioned by their p-values from an impact analysis measuring total perturbation accumulation (pAcc) vs. a classic over-representation analysis (pORA). Pathways with FDR p-value < 0.05 by both analyses are shown in red. Selected pathways are highlighted in yellow.

### An orthogonal model system confirms a canonical tubular response to serum-induced injury

Next, we sought to confirm our findings in an orthogonal and supraphysiologic system by comparing the transcriptional and epigenomic response of primary human tubular epithelial cells cultured in 10% FBS to intact renal cortex. Comparison of cultured human tubules to intact renal cortex revealed 939 genes that were also differentially expressed in the tubule MPS system. Of these, 661 (70.4%) also showed significant differential expression with the same directionality i.e., up- or down-regulated as the MPS system (**Figure 3A**). Gene ontology analysis of all differentially expressed genes identified significant enrichment of biological processes related to cellular proliferation in upregulated genes while solute transport processes were enriched in the downregulated genes (**Figure 3B**). The overall pattern of increased expression of genes related to cellular proliferation, extracellular matrix reorganization and inflammatory cytokine secretion with downregulation of genes related to metabolism and solute transport was similar to that seen for tubule MPS (**Figure 3C**, compare to **Figure 2C**). These overall gene ontology enrichments were also very similar to a standard model of acute kidney injury, the murine unilateral ureteral obstruction model (**Supplemental Figure 1**) (*36*). In Advaita iPathwayGuide analysis, metabolic (p=2.150×10^−21^) and cell cycle pathways (p=4.796×10^−6^) were significantly enriched as were cytokine/chemokine-mediated signaling pathways (p=3.623×10^−5^), similar to that seen with serum-treated tubule MPS (**Figure 3D**).

**Figure 3.**
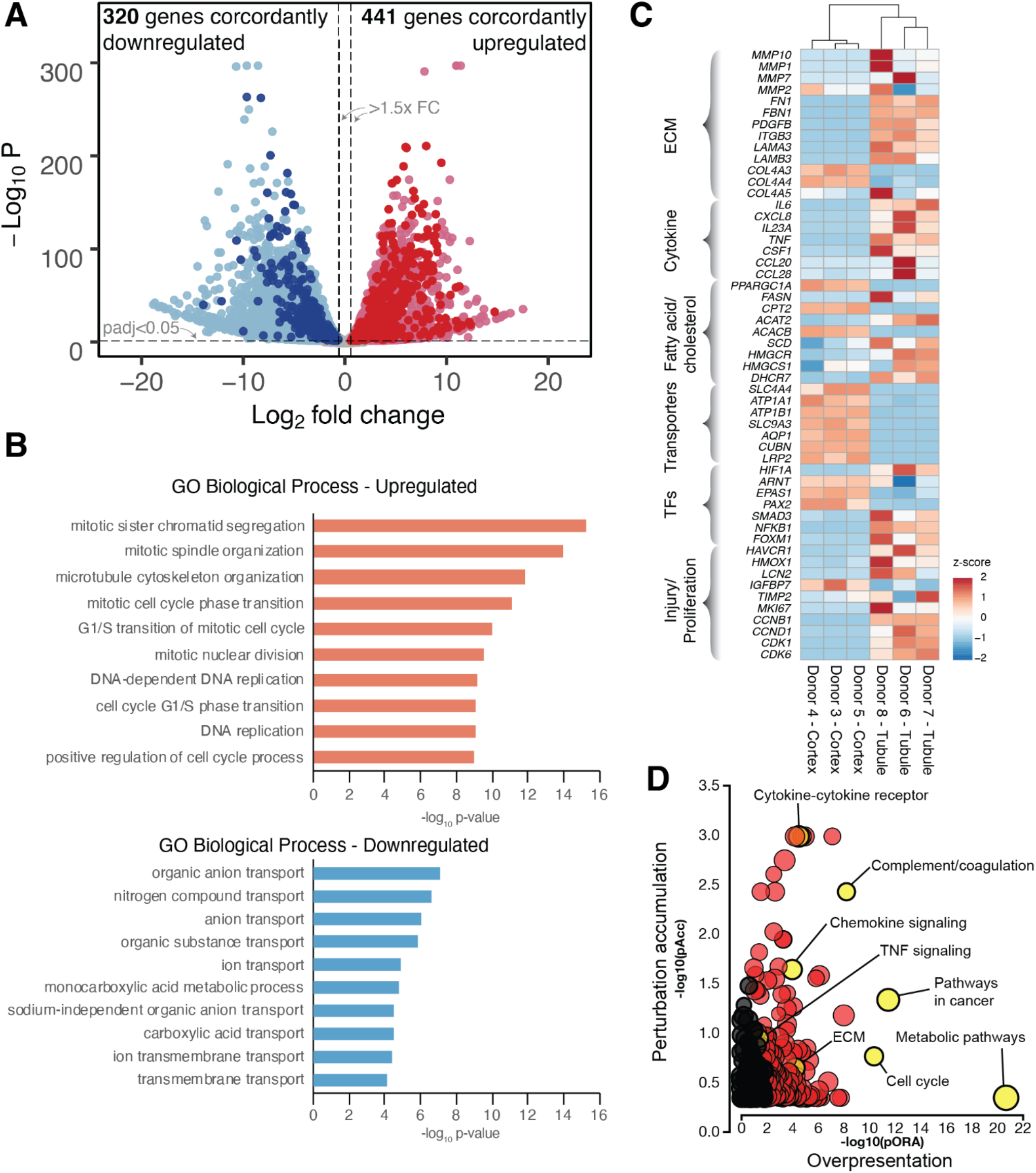
Global transcriptional changes induced in primary tubules cultures in 10% FBS. A) Volcano plot of differentially expressed genes between intact renal cortex and kidney tubules cultured in 10% FBS (lighter dots). Upregulated = higher expression in cultured tubules. The darker dots represent genes that exhibit the same directionality of significant gene expression change in the tubular MPS serum exposure experiment (Figure 2A). B) Gene ontology enrichments for top 10 up- and down-regulated biological process in the differentially expressed gene set. C) Heatmap of the same exemplar genes as shown in Figure 2C showing a consistent pattern of expression changes across the two systems. D) Advaita pathway analysis demonstrates significant perturbation of metabolic, cytokine/chemokine, and cell cycle pathway components in serum-exposed tubule cells. Dots representing pathways are positioned by their p-values from an impact analysis measuring total perturbation accumulation (pAcc) vs. a classic over-representation analysis (pORA). Pathways with FDR p-value < 0.05 by both analyses are shown in red. Selected pathways are highlighted in yellow.

### Distinct epigenomic landscape of adult human kidney cortex

To gain insight into the regulatory processes driving the observed transcriptional differences, we generated chromatin accessibility profiles from intact renal cortex and medulla from 3 donors using ATAC-seq. This identified 83,124 regulatory elements (**Supplementary Table 1**) in intact kidney parenchyma, only 22.3% of which were within 5kb of known gene transcription start sites (TSS) (**Supplemental Figure 2A**). This is in contrast to a recent single nucleus ATAC-seq study of human kidney, in which >50% of the identified regulatory elements are within an even narrower window of <3kb from known TSS (bioRxiv 2020.06.14.151167; doi: https://doi.org/10.1101/2020.06.14.151167). This difference may reflect increased sensitivities or efficiencies of library construction of bulk versus single-nucleus chromatin profiling approaches. We have previously shown that genetic variants linked to kidney disease and traits are enriched in kidney cell type-specific regulatory regions. Using our newly generated chromatin accessibility maps, we localized 54 kidney disease associated GWAS loci to regulatory elements seen only in intact adult kidney cortex (**Supplementary Table 2**). This included two additional loci (rs4293393, rs12917707) which mapped to open chromatin regions in the *UMOD* locus (**Supplemental Figure 2B**). In addition, 1,883 of the regulatory elements from intact kidney cortex were not detected in the most recent and comprehensive regulatory element index generated from 733 different cell and tissue samples by ENCODE and were therefore unique to intact adult kidney tissue (*37*). Notably, this ENCODE index included cultured tubular epithelial cells, primary glomerular cultures, renal cell carcinomas and fetal kidney samples but did not contain intact adult kidney tissues. 95.6% of these 1,883 adult kidney unique regulatory elements were located >5kb from known TSS, characteristic of distal regulatory elements such as enhancers (**Figure 4A**). GREAT analysis revealed that these elements were located close to genes whose molecular function ontologies described potassium channels and transporter activity (**Figure 4B**). The majority of these genes had lower expression levels in tubules cultured in 10% FBS compared to intact renal cortex (**Figure 4C**). Conversely, metabolism related genes (*GAPDH, ME2, HMGCR*) had higher expression in the cultured tubule samples. Restricting our analysis to adult kidney derived primary cultures, cell lines and intact kidney samples, we generated a master list of 555,884 regulatory elements. Within this, 44,756 elements (8.1%) showed significantly higher accessibility in tubules cultured in 10% FBS compared to intact renal cortex (**Figure 4D**). Conversely, 37,053 elements (6.7%) showed higher accessibility in intact renal cortex compared to cultured tubules. These differentially accessible regulatory elements represented the portion of the epigenome that was driving gene expression differences between these sample types (**Supplementary Table 3**). More generally, differentially accessible regulatory elements were more likely to be located near genes exhibiting differential expression between cultured tubules and intact cortex (**Supplementary Figure 2C**). For example, 2,941/11,633 (25.3%) of differentially expressed genes had a differentially accessible regulatory element within 2.5kb of their TSS. In contrast 5,828/49,017 (11.9%) of non-changing genes had at least one differentially accessible regulatory element near their TSS (Chi-squared with Yates Correction p<0.0001). GREAT analysis of the elements with greater accessibility in intact renal cortex revealed localization to genes with molecular functions associated with transporter and transcription factor activities (**Figure 4E**). Several of these transcription factors, *PPARA, HNF4A, VDR, RXRA, RARA* and *ESRRA* demonstrated increased chromatin accessibility in renal cortex that was localized both to their TSS and at distal sites and this corresponded with increased gene expression in intact renal cortex (**Figure 4F** and **Supplementary Figure 3**). PPARγ coactivator-1α (PGC-1α), a major regulator of mitochondrial biogenesis and metabolism and interaction partner of PPARγ and ESRRA displayed 1.9x and 13.3x reduced expression in the MPS and 2D culture systems, respectively.

**Figure 4.**
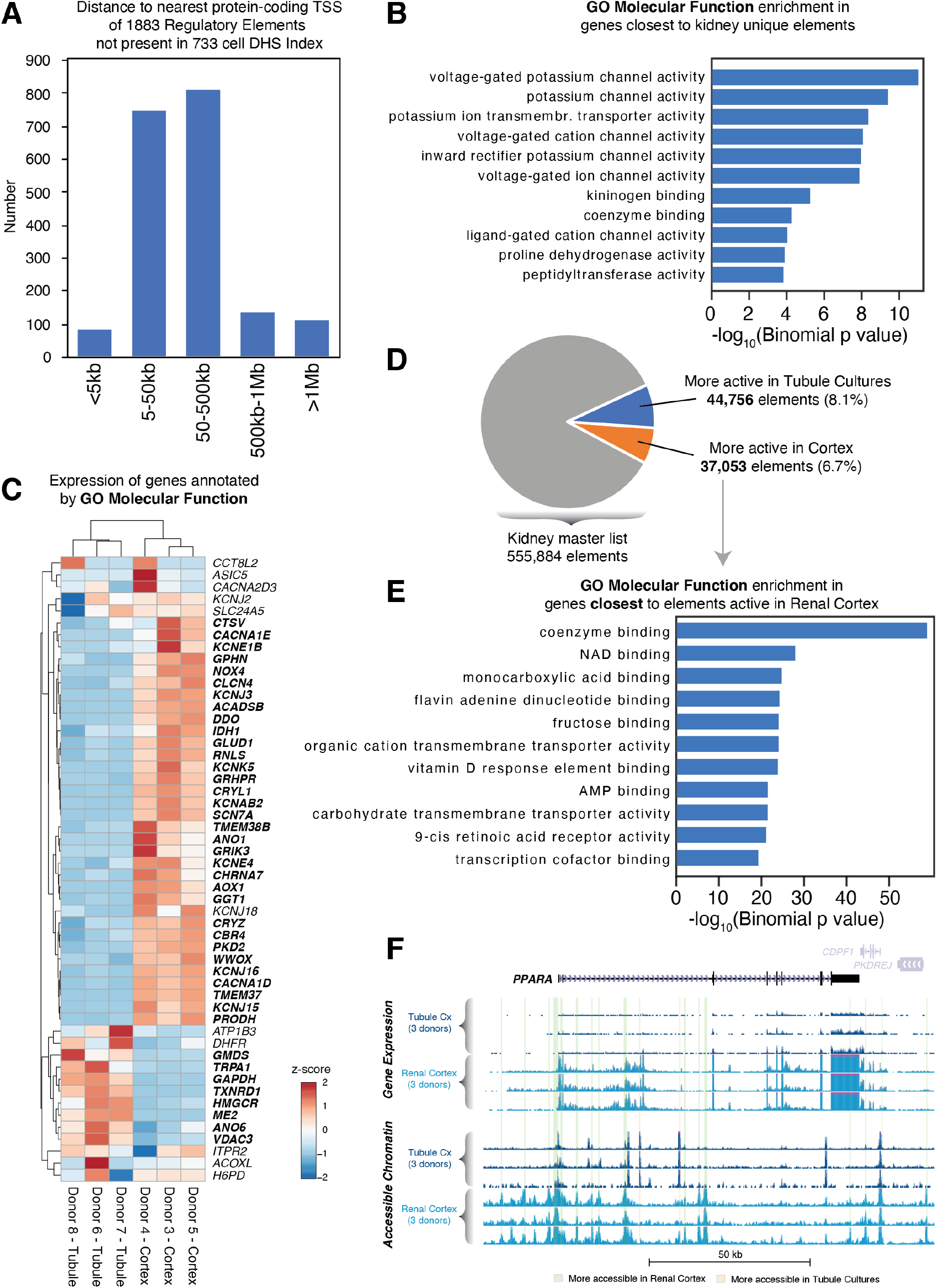
The distinct regulatory landscape of adult human kidney. A) Distribution of the distance to the nearest gene’s TSS for the 1,883 regulatory elements that are only present in accessible chromatin of adult human kidney and not in the 733-cell ENCODE regulatory element index. B) Stanford GREAT analysis GO enrichment of molecular function of the nearest genes associated with the adult kidney restricted regulatory elements. C) Heatmap of all the genes associated with the molecular function terms shown in panel B. Genes in boldface show significantly different expression between intact kidney cortex and tubules cultured in 10% FBS. D) Proportion of regulatory elements with significantly higher accessibility in either renal cortex or cultured tubules. E) Stanford GREAT analysis GO enrichment of molecular function of the nearest genes associated with regulatory elements showing greater accessibility in intact renal cortex compared to cultured tubules. F) Overview of the *PPARA* gene locus with gene expression (top) and accessible chromatin (bottom) tracks for cultured tubules and intact renal cortex (3 donors each). The PPARA gene is highly expressed in intact renal cortex but not in cultured tubules. This corresponds to numerous regulatory elements (vertical green bars) that show greater accessibility in intact cortex compared to cultured tubules.

### Transcription factor drivers of serum-induced tubular transcriptional response

Next, we used two complementary approaches to understand the influence of transcription factors on gene regulation. First, we used ENRICHR which seeks to understand the impact of transcription factors by examining the expression changes of their known target genes annotated by ChIP-seq (i.e., ChEA database). ENRICHR-ChEA implicated roles for NF-κB, SMAD3, AP-1 and FOXM1 transcription factors in orchestrating the tubular response to serum exposure in the both the MPS and tubule/intact cortex systems. Many of these transcription factors had increased expression in serum-exposed tubules in both of our model systems (**Figure 5A, B**). The lack of significant induction of *JUN*, an AP-1 component is consistent with its known regulation at the post-transcriptional level. In a complementary approach, we leveraged our chromatin accessibility datasets to ask which transcription factor binding motifs were enriched in regulatory elements with greater accessibility in intact renal cortex vs. tubules cultured in 10% FBS. Using HOMER, we found that regulatory elements selectively accessible in renal cortex were highly enriched in binding motifs for HNF4A, PPARA, PPARG, RARA, ESRRA and RXR transcription factor families (**Figure 5C**). Conversely, the regulatory elements with greater accessibility in cultured tubules exhibited a strong enrichment for AP-1, NF-κB and FOXM1 transcription factor binding motifs (**Figure 5D**), some of which also exhibit increased expression in serum-exposed tubules (**Supplemental Figure 4**). Taken together, the complementary and orthogonal analyses (GREAT, ENRICHR, HOMER) depicted in **Figures 5 and 6** demonstrate that serum-exposed tubules exhibit a strong stress-response signature coordinated by AP-1, NF-κB and FOXM1 transcription factors.

**Figure 5.**
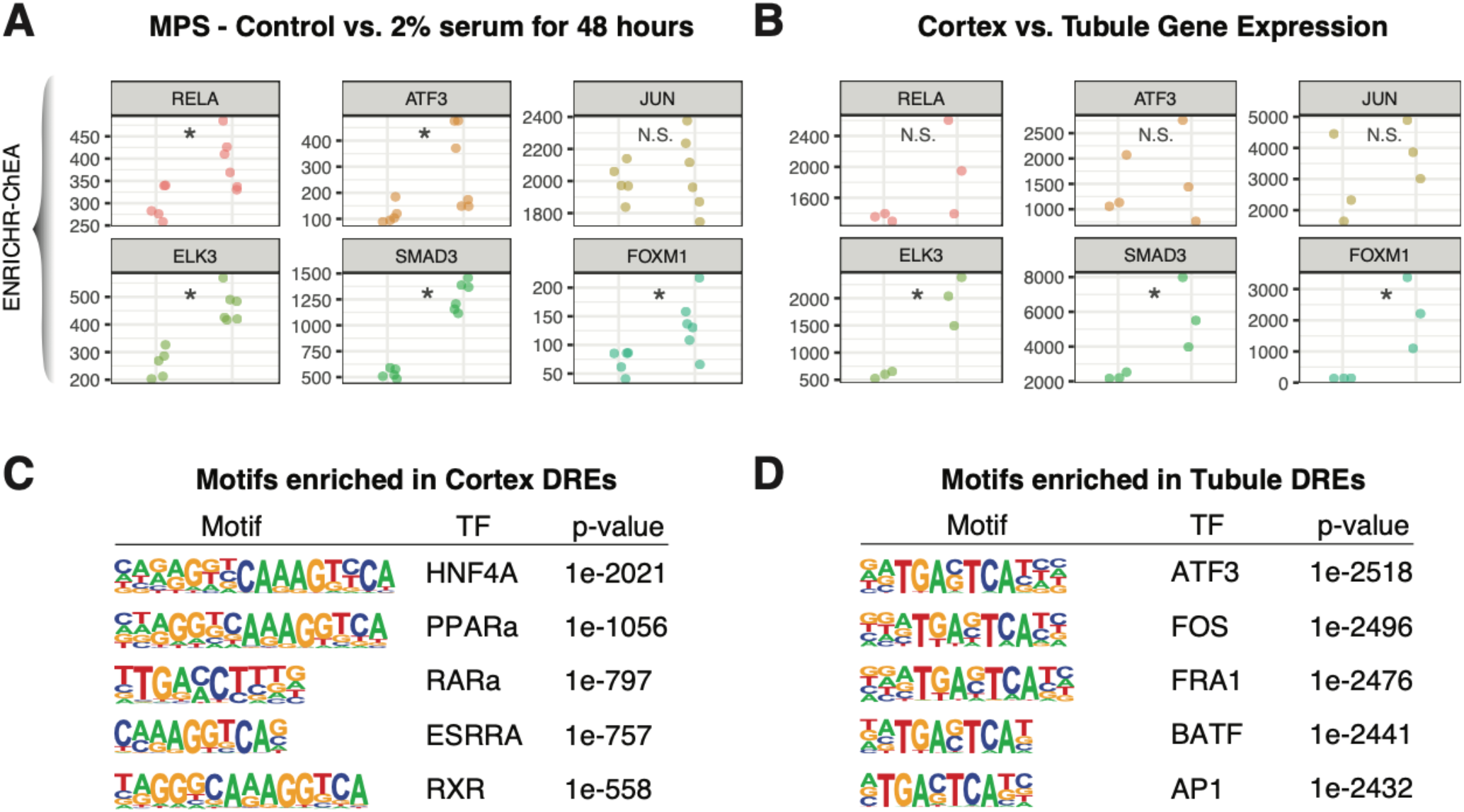
Transcription factors drivers of the transcriptional landscape of injured tubules. Gene expression dot plots of transcription factors implicated by ENRICHR-ChEA as having significant changes in expression of their known target genes in A) tubule MPS and B) tubules cultured in 10% FBS. Transcription factor binding motif enrichments in regulatory elements with higher accessibility in C) intact renal cortex and D) tubules cultured in 10% FBS.

**Figure 6.**
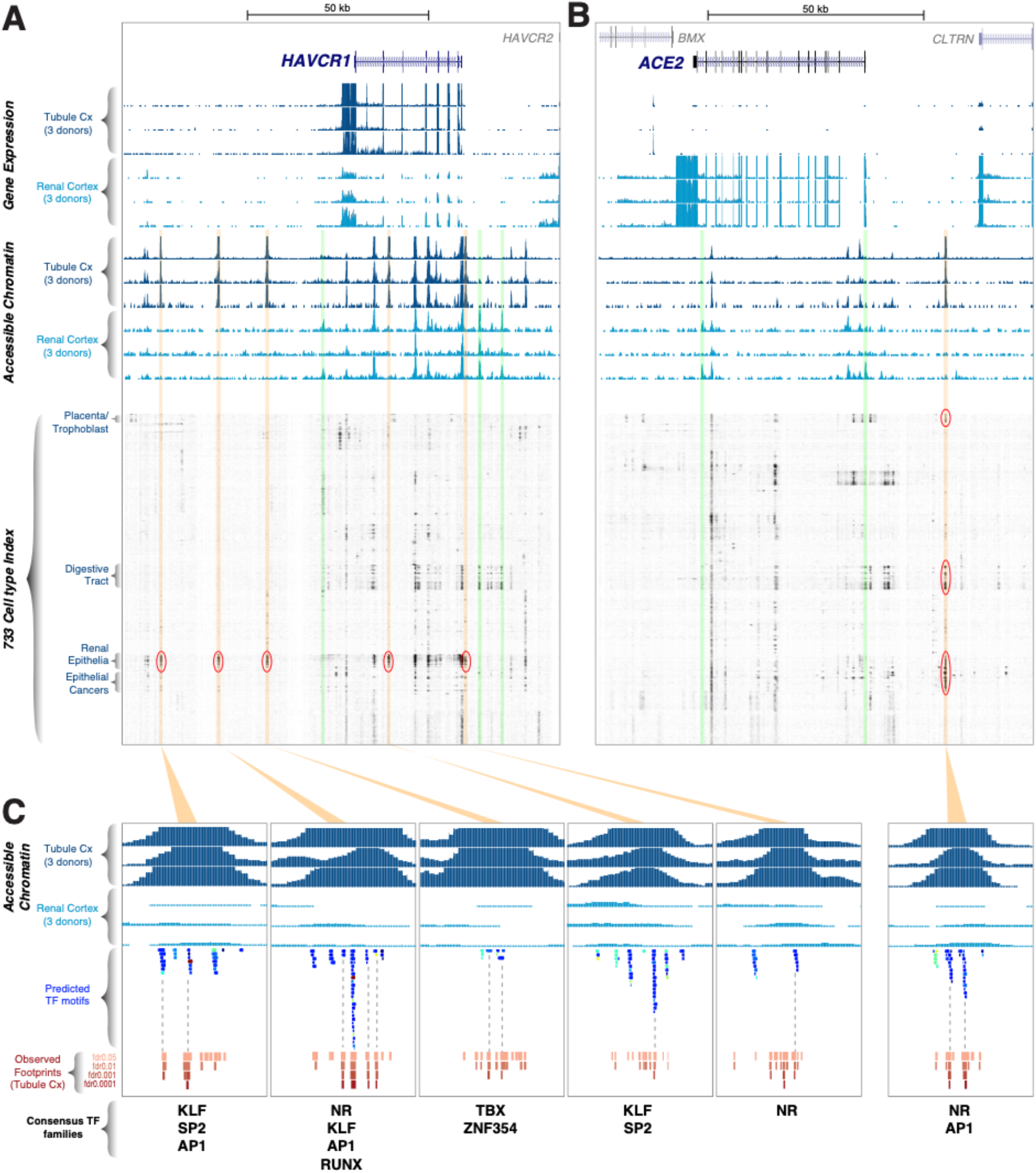
Integrative analysis reveals unique regulatory circuitry of *HAVCR1* and *ACE2*. Overview of the A) *HAVCR1* and B) *ACE2* gene loci with gene expression (top) and accessible chromatin (middle) tracks for cultured tubules and intact renal cortex (3 donors each). The *HAVCR1* gene is expressed at higher levels in tubules cultured in 10% FBS whereas *ACE2* shows the opposite pattern of expression. Green vertical bars indicate regulatory elements with greater accessibility in intact cortex, while orange vertical bars indicate regulatory elements with greater accessibility in cultured tubules. Lower panel shows the activity of the same loci in the 733-cell ENCODE regulatory index. Elements with selective activity patterns around both genes are highlighted. C) Comparison of the predicted transcription factor motif archetypes to actual footprints (FDR≤0.001) in tubule DNase-seq data reveals overlap of predicted motifs for AP1 and NR families in kidney-selective regulatory elements identified in panels A and B.

### Integrative analysis of kidney injury and COVID-19 associated gene loci

We then performed in depth integrated analysis to demonstrate the utility of our datasets and derive new understanding of the regulation of exemplar gene loci. Regulatory elements can exhibit extremely cell type and state-dependent patterns of accessibility and therefore, we were aided by the recent publication of the ENCODE regulatory element index derived from 733 cells (*37*). This index permitted identification of elements restricted to particular cell types or lineages, but in our particular instance was biased toward detection of elements in cultured tubule cells (i.e., missing elements that would only be seen in intact renal cortex, as described in **Figure 5**). Using deeply sequenced chromatin accessibility profiling data, the nucleotide sequence bound by a transcription factor appears as a region of protection (a footprint) in the broader region of open chromatin associated with that regulatory element. By reading the underlying DNA sequence and matching it against transcription factor motif archetypes, it is possible to infer transcription factor binding at a particular genomic location (at least at the level of transcription factor family with shared binding motifs). Recently, a large-scale effort to map transcription factor motif archetypes and actual footprints in multiple cell types/states has been described (*38*). First, we integrated these two recently published datasets together with our data to study the classic kidney injury biomarker gene *HAVCR1* (KIM1). The *HAVCR1* gene is expressed at 23x higher levels in cultured tubular epithelial cells compared to intact renal cortex. This increased expression corresponds to open chromatin, with at least 5 regulatory elements in cultured tubules (vertical orange bars) that were not accessible in intact renal cortex (**Figure 6A**). In the 733-cell index, these regulatory elements exhibit an activity profile restricted to renal epithelial cells suggesting a kidney-specific role in *HAVCR1* gene regulation. Other kidney injury marker genes including *LCN2* (increased 64x), *HMOX1* (increased 4.3x) and *QPRT* (decreased 3.4x) also showed distinct differentially accessible regulatory elements, though not all of these showed kidney-restricted patterns in the 733-cell regulatory index (**Supplemental Figure 5A-C**).

Acute tubular injury is a frequent finding in patients with COVID-19 and so we asked if our models of tubular injury could provide insight into the regulation of COVID-19 associated genes. First, we explored the regulatory landscape around the *ACE2* gene, which is highly expressed in proximal tubules and is a known entry receptor for SARS-CoV-2, the causative agent of COVID-19 disease. *ACE2* is highly expressed in intact renal cortex, but its levels are 385x lower in cultured tubules. A singular regulatory element shows greater accessibility in cultured tubules and its anti-correlation with *ACE2* expression suggests a potential repressor function (**Figure 6B**, vertical orange bar). High resolution examination of these regulatory elements around *HAVCR1* and *ACE2* revealed overlap of predicted transcription factor motif archetypes with actual footprints in kidney tubule cells even at increasingly stringent FDRs (**Figure 6C**). This analysis implicates KLF, AP1 and nuclear receptor (NR) transcription factor families in the regulation of *HAVCR1* and *ACE2* gene expression. The presence of AP1 and NR binding motifs and footprints in putative activating (*HAVCR1*) and repressive (*ACE2*) regulatory elements is consistent with their pleomorphic roles in activation and repression of gene transcription in different contexts. Other genes associated with SARS-CoV-2 infection such as *TMPRSS2* (decreased 21x), *IL6* (increased 2,435x) and *CXCL8* (increased 194x) also showed numerous differentially accessible regulatory elements at their genomic loci (**Supplemental Figure 5D-F**).

### Serum exposed MPS excrete inflammatory cytokines and matrix metalloproteinases into their effluents

To confirm the canonical stress response of tubular epithelial cells predicted from our transcriptomic and epigenomic studies, we measured protein levels of pro-inflammatory cytokines (IL6 and CXCL8), matrix remodeling enzymes (MMP1 and MMP7) and shed HAVCR1 in serum exposed MPS effluents from 3 additional donors by ELISA (**Figure 7**). We collected effluents from early (8 hours) to later timepoints (7 days) and also exposed a parallel set of MPS from the same donors to purified human albumin at equivalent concentration present in 2% human serum (720 μg/ml). For the majority of conditions, serum treatment rapidly and significantly increased secretion of these analytes relative to control, confirming the predictions from our multi-omic studies. In contrast, albumin treatment induced only modest secretion of KIM-1 or MMP7 in 1-2 donors. To confirm these findings in these additional donor samples, we performed transcriptomic analysis of serum or albumin treated MPS at 7 days. Overall, both 2% human serum and albumin suppressed the expression of genes related to fatty acid oxidation, TCA cycle and mitochondrial electron transport chain, and ion transport. By contrast, only serum treatment induced genes related to ECM reorganization, inflammation, and cell stress and proliferation (**Supplemental figure 6**). These results orthogonally validated our multi-omic findings and demonstrated the power of MPS systems for dissection of tubular injury responses to defined stimuli.

**Figure 7.**
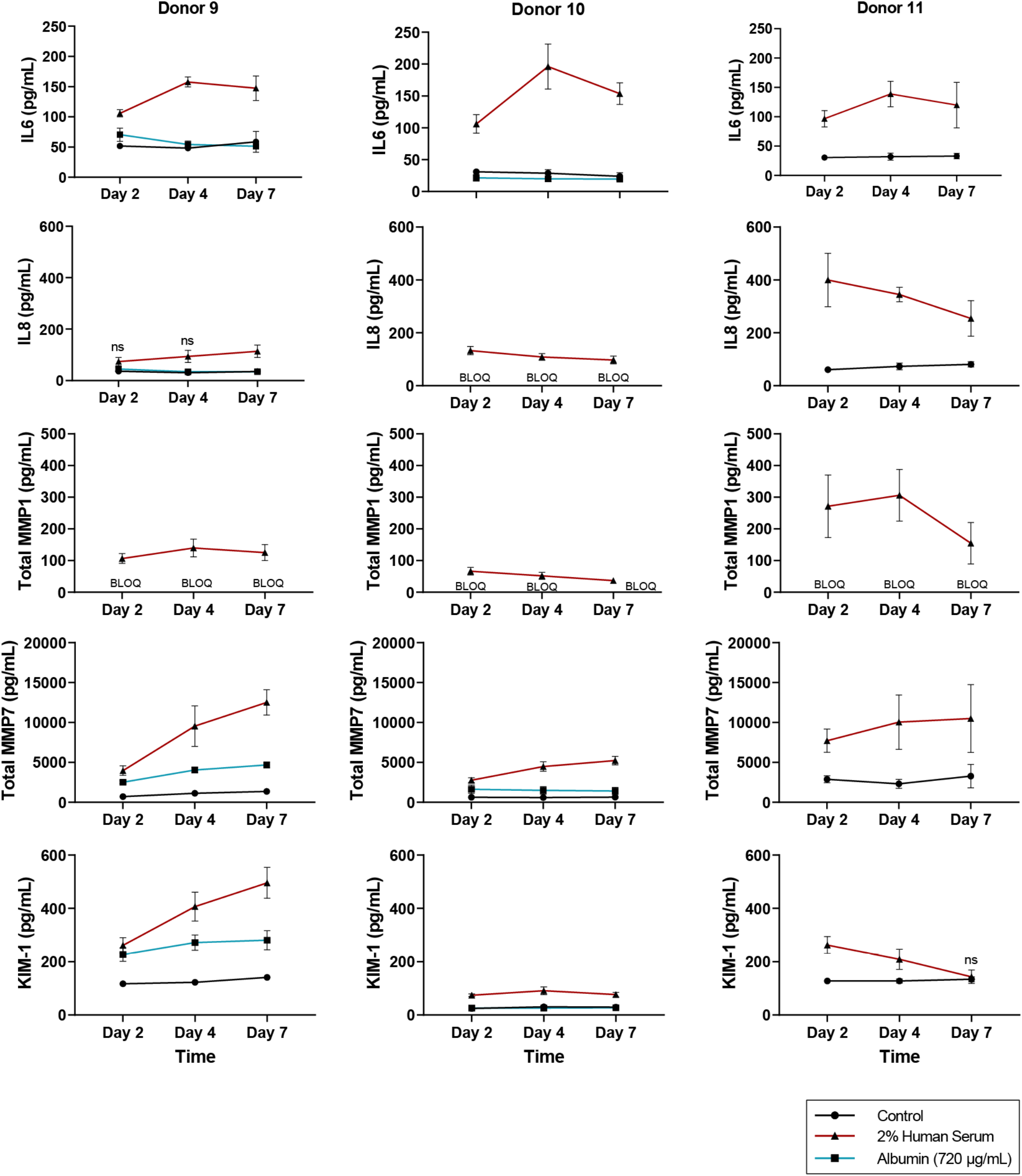
Serum exposure induces secretion of cytokines, matrix metalloproteinases and shedding of HAVCR1 in 3D MPS. Effluents from serum and albumin treated 3D MPS from 3 donors were assayed for the indicated proteins by ELISA. All comparisons included at least 4 replicates for each timepoint and treatment, except for Donor 9, for which only 1 replicate was available for the day 2 and day 4 timepoints. Except where indicated as not significant (ns), all comparisons were significant compared to the control sample at an adjusted p-value < 0.05 using an unpaired t-test with Sidak-Holm correction for multiple comparisons. BLOQ, below limit of quantification.

## Discussion

Here we demonstrate that exposure of human kidney tubular epithelial cells induces proliferation and upregulated transcription of TNF-signaling associated cytokine and ECM remodeling genes. Concomitantly, there is downregulation of genes for transporters and metabolic and biosynthetic programs. This coordinated response is consistent across both 2D and 3D model systems of tubular injury and is similar to pathways that are activated in the murine UUO model of kidney injury (*36*). To derive the mechanisms driving these changes, we generated high-resolution chromatin accessibility profiles of intact human kidney cortex. We found that this baseline epigenomic state is characterized by hundreds of distinct regulatory elements with features of nuclear receptor activity from several families (HNF4A, PPARA, ESRRA, RXRA, RARA, VDR). By contrast, the tubular response to serum exposure is coordinated by stress-activated transcription factors in the AP-1 and NF-κB families. Regulatory elements around the kidney injury biomarker gene *HAVCR1* display a kidney-restricted pattern of accessibility and contain footprints compatible with binding of specific transcription factor families. We identified similar switch-like regulatory elements around other kidney injury biomarker genes and around genes associated with COVID-19 disease. Lastly, we confirmed our findings by demonstrating secretion of cytokines and matrix remodeling enzymes in effluents of 3D tubule cultures exposed to serum.

Our studies establish that a key feature of the tubular epithelial cell response to injury, irrespective of the nature of the inciting stimulus, is downregulation of oxidative phosphorylation, transporters and channels. Both serum and albumin induced this injury response, though only serum also activated a proinflammatory secretory response as well (see below). PPAR-mediated signaling plays a central role in oxidative phosphorylation in metabolically active tissues (*39*). Therefore, the loss of open chromatin regions containing PPAR-family binding motifs together with decreased expression of their known transcriptional coactivator, *PPARGC1A* (encoding PGC-1α) is consistent with the idea that expression and function of PPAR-family transcription factors is reduced in injured tubules. PGC-1αlevels are also reduced in animal models of sepsis- and folic acid-induced AKI (*40, 41*) and in human AKI patient biopsies (*42*). Reduced PGC-1αactivity together with loss of the rate-limiting enzyme QPRT, whose gene expression we observed is also reduced in injured tubules, can lead to lower levels of niacinamide predisposing to kidney injury (*42, 43*). The loss of HNF4A expression and its inferred chromatin binding in injured tubules is also consistent with its role in regulating the expression of the urate transporter ABCG2 (*44*), the Na+/H+ antiporter 3, NHE3 (*SLC9A3*) (*45, 46*) and the electrogenic Na+/HCO3 cotransporter, NBC1 (*SLC4A4*). The distinct epigenomic landscape of intact adult human kidney also revealed substantial regulation of numerous potassium channel genes (*KCNJ3, KCNK5, KCNAB2, KCNE4, KCNJ15* and *KCNJ16*; **Figure 5**). While many of these channels appear to be expressed in kidney cortex (ProteinAtlas), the function of most remains to be elucidated. One exception is the *KCNK5* gene, whose gene product is also known as TASK2 and whose crystal structure was recently solved (*47*). This two-pore domain, acid-sensitive potassium channel is expressed in the proximal tubule and collecting ducts of the kidney (*48, 49*). TASK2 plays a key role in bicarbonate reabsorption in the proximal tubule and its deletion in mice results in defects in renal proximal tubular acidosis (*49*). Interestingly, missense mutations in *KCNK5* have been identified in a small cohort of patients with the chronic tubulointerstitial disease Balkan endemic nephropathy (*50*).

We also demonstrate that serum, but not albumin-injured tubules secrete pro-inflammatory cytokines and matrix remodeling enzymes. TNFαcan be produced by tubular epithelial cells (*51*) and its levels are increased during injury due to active rejection (*52*), cisplatin (*51, 53*) or unilateral ureteral obstruction (*54*). After serum exposure, we were able to detect IL6 and CXCL8 in MPS effluents which appears to mimic the increased levels of those same cytokines in urine from patients with ischemic injured allograft kidneys (*55*). We also detected secreted MMP7 and MMP1 within MPS effluents following injury. Urinary MMP7 has recently been described as an adverse prognostic biomarker of AKI (*56, 57*). MMP7 degrades E-cadherin and can release β-catenin (*58*) which in turn can reinforce MMP7 expression (*59*). MMP7 is also induced in folic acid nephropathy and UUO animal models (*60*). Besides being a biomarker, MMP7 may in fact be a pathogenic driver of kidney fibrosis (*56*). Diabetic kidney disease is the most common etiology for proteinuria and in proteinuric diabetic patients, elevated urinary MMP7 levels are associated with progressive kidney disease and increased mortality (*61*). In contrast, relatively little is known about the role of MMP1 in AKI. Therefore, further investigation of the role of MMP1 in AKI and subsequent remodeling of the tubulointerstitium is warranted.

The exact concentration of albumin that is physiologically present in the proximal tubule is a matter of continuing debate. It is thought to vary from ~30 μg/ml (glomerular sieving coefficient for albumin, GSC_A_ ~ 0.00062) determined from early micropuncture studies in the rat (*62–65*) to almost 40-50x higher, ~1.2 mg/ml (GSC_A_ ~ 0.025) using more recent two photon microscopy based approaches (*66, 67*). The concentration of albumin used in our study (720 μg/ml) falls between these estimates. This amount of albumin if delivered to the final urine without reabsorption would result in albumin excretion of >1g/24 hrs. Pure albuminuria (selective proteinuria) is typical of minimal change disease (*68*) which is associated with good prognosis and little if any residual tubulointerstitial fibrosis. In contrast, non-selective proteinuria with albumin and other serum proteins being delivered to the tubules is seen with diseases such as focal and segmental glomerulosclerosis and diabetic nephropathy which can be associated with progressive tubulointerstitial injury and fibrosis (*69, 70*). Of course, albumin can bind to numerous molecules whose composition and abundance in the serum can vary between healthy and disease states. Albumin is also subject to various modifications (*71*) that can alter its biodistribution and pharmacodynamics as was recently demonstrated for carbamylated albumin (*72*). Our studies demonstrate that normal purified albumin alone does not induce a proinflammatory secretory phenotype for which other components of the serum appear necessary. In future studies, multi-omic characterization of MPS exposed to patient derived albumin (e.g. glycated forms) (*73*) or albumin ± other serum components will prove useful to dissect tubular injury mechanisms.

Bulk epigenomic profiling methods such as we have utilized in our study are ideally suited for comprehensive and deep mapping of cellular phenotypes and complement the granularity of single cell-based approaches. Therefore, we compared our findings to those of two recent single nucleus and single cell-based profiling studies of human and mouse kidney. Muto and colleagues integrated single nucleus RNA-seq and ATAC-seq to identify cell-type heterogeneity within the human kidney (bioRxiv 2020.06.14.151167; doi: https://doi.org/10.1101/2020.06.14.151167). Park and colleagues used single cell RNA-seq and ATAC-seq as well as bulk RNA-seq to characterize cell-specific changes in human and mouse kidney during disease states (bioRxiv 2020.09.21.307231v1; doi: https://doi.org/10.1101/2020.09.21.307231). Overall, our data presented here is in good agreement with their findings – namely that HNF4A, PPARA, and ESSRA transcription factor families are important for maintenance of cell identity and control of expression of genes related to cellular metabolism and transport. In addition, the activity of stress-associated transcription factors such as NF-κB are enriched in PTEC subpopulations during injury, which may play a role in acquisition of a proinflammatory gene signature. Indeed, NF-κB has previously been implicated in driving the inflammatory response of PTECs to high molecular weight protein challenge, and its inhibition has been shown to mitigate pro-inflammatory cytokine production and prevent renal fibrosis in animal models of kidney disease (*19, 74, 75*). Arrest of proximal tubule cells in G1/S or G2/M phases of the cell cycle after injury has also been proposed as a mechanism by which PTECs acquire a senescence associated secretory phenotype (*25, 76*). We do not observe a prominent transcriptional signature consistent with G2/M arrest, such as induction of cyclin G1, p53, or other DNA damage response transcripts. However, it is possible that a subpopulation of the cells did undergo G2/M arrest, but this signature was masked due to using a bulk RNA-seq approach. On the other hand, serum treatment caused secretion of KIM-1 and enrichment of FOXM1 epigenomic motifs, events that are consistent with cell dedifferentiation and proliferation in tubular wound healing (*23*).

Our data also provide insights into the regulation of ACE2, the best studied entry receptor for SARS-CoV-2, the causative agent of COVID-19 (*77*). *ACE2* is highly expressed in proximal tubular epithelial cells (*78*) and blockade of this entry pathway can prevent viral infection of kidney organoids (*79*). Acute kidney injury is common in COVID-19 patients (*80*) and this has raised the question whether this is mediated by direct viral infection of the kidney via ACE2. However, while direct infection of the kidney parenchyma may be anecdotally possible, the majority of biopsy and autopsy studies to date do not support sustained and significant infection of the kidney by SARS-CoV-2. The best consensus is that kidney injury in COVID-19 patients is due to pre-renal mechanisms related to pulmonary dysfunction and/or systemic derangements due to secretion of inflammatory cytokines (*81*) rather than direct kidney infection by the virus. Here, we show that canonical stress-inducible transcription factors control the overall tubular transcriptional response to injury and that *ACE2* is downregulated with this treatment. Our finding is consistent with previous studies showing that protein overload reduces ACE2 gene expression via an NF-κB pathway in unilaterally nephrectomized rats (*82*) and in HK-2, a human proximal tubular epithelial cell line (*83*). By counteracting the effects of angiotensin I and II, angiotensin 1-7 and 1-9, the products of ACE2 activity are thought to be kidney protective (*84*). Therefore, in COVID-19 downregulation of ACE2 in the setting of tubular injury may ultimately be maladaptive for the kidney and promote progression of CKD.

One limitation of our study is the limited number of samples, which is somewhat mitigated by consistent trends across biological replicates. A major advantage is our use of primary human kidney cells and tissues and orthogonal validation in the MPS. Altogether, the data from the PTEC-MPS are in good concordance with previous observations that proteinuria can drive the proximal tubule to acquire a proinflammatory signature, suggesting that it may be a suitable human-relevant model for evaluating novel approaches to impede tubular acquisition of a maladaptive phenotype. Furthermore, our chromatin accessibility datasets provide a deep reference map of the unique regulatory architecture of intact human kidney and its response to serum-induced injury which will inform future GWAS of kidney injury and emerging triggers of acute kidney injury such as COVID-19.

## Methods

### Human kidney tissues and primary cultures

Samples were collected in deidentified fashion through Northwest Biotrust at the University of Washington Medical Center with local IRB approval (Study 1297). Donor demographics are listed in **Supplementary Table 1**. Both primary tubular cell cultures and intact renal tissue were derived from tumor-adjacent normal tissue in nephrectomy specimens. For intact tissues, macrodissected portions (~100-200mg) of renal cortex and medulla were snap frozen in liquid nitrogen and stored at −80°C until experiments were performed. Aliquots were also stabilized in RNALater (Invitrogen) overnight at 4°C before long term storage at −80°C. Isolation and culture of primary tubular epithelial cells in the presence of 10% FBS has been described earlier (*85*). Briefly, after protease digestion, cortex fragments were placed in tissue culture treated flasks enabling outgrowth of cortical tubular epithelial cells. Cells were grown in RPMI supplemented with 10% FBS and insulin-transferrin-selenite+ [ITS+] supplement. Isolation and culture of primary tubular epithelial cells in serum-free media has also been previously described (*30, 86*). Serum-free tubular cell cultures were maintained in DMEM/F12 (Gibco, 11330-032) supplemented with 1x ITS-A (Gibco, 51300044), 50 nM hydrocortisone (Sigma, H6909), and 1x Antibiotic-Antimycotic (Gibco, 15240062). For passaging, the cells were subjected trypsin digestion using 0.05% trypsin EDTA (Gibco, 25200056) and manual cell scraping upon reaching 70-80% confluence. The single-cell suspension solution was neutralized with defined trypsin inhibitor (Gibco, R007100), centrifuged at 200 × g for 5 mins, and resuspended in maintenance media. All experiments were performed with cells passage 2-4.

### Microphysiological devices

Triplex™ microfluidic devices were purchased from Nortis, Inc and prepared as previously described with slight modification (*28, 86*). Briefly, device chambers were filled with 6 mg/mL rat tail type I collagen (Corning, 354236) and the matrix was allowed to polymerize overnight at room temperature. Microfiber inserts were removed, and the resultant channel was coated by injecting 2 μL of 0.1 mg/mL collagen IV (Corning, 354233) using a 5 μL syringe (Hamilton, 7634-01) outfitted with a 22-gauge small hub needle (Hamilton, 7804-01). Devices were incubated for 30 minutes at 37 degrees Celsius before initiating flow at 1 μL/min and equilibrating the system with maintenance media for 2 hours. PTECs were harvested from culture vessels and resuspended at a concentration of ~20 × 10^6^ cells/mL, and ~2.5 μL was injected into each channel. The cells were allowed to adhere for 2-4 hours before initiating flow at 1 μL/min. During treatment, effluent was collected from outflow collection reservoirs at specified timepoints and transferred to microcentrifuge tubes for storage at −80°C until analysis.

### Immunofluorescence microscopy

All procedures were performed at room temperature with a flow rate of 10 μL/min for all solutions. At the end of treatment, the cells were fixed by perfusing 10% phosphate buffered formalin (Fisher, SF100-4) for 30 minutes followed by a 60-minute wash with dPBS (ThermoFisher, 14040133). The devices were stored at 4 degrees Celsius for not more than 2 weeks before further processing. To prepare the tubules for labeling of Ki-67, the channels were first blocked and permeabilized with dPBS containing 5% bovine serum albumin (Sigma, A2153) and 0.1% Triton X-100 for 2 hours. Rabbit monoclonal anti-Ki67 (Abcam, ab16667) was diluted 1:10 in dPBS+5% BSA and 10 μL was injected into each channel and allowed to incubate for 1 hour. The channels were washed with dPBS+0.05% tween-20 for 2 hours followed by a 1-hour perfusion of goat anti-rabbit secondary (Fisher, A11037) diluted 1:1000 in dPBS+5% BSA. The channel was washed once more with dPBS+0.05% tween-20 for 2 hours before the nuclei were labeled by perfusion of 1 ug/mL Hoechst 33342 in dPBS for 30 minutes followed by a 30-minute dPBS wash. The cells were imaged on a Nikon Eclipse Ti-S microscope equipped with a Nikon DS-Fi3 camera. The total number of nuclei and Ki67 positive nuclei in each 10x image were determined by manual count with the multi-point tool in ImageJ. The percentage of nuclei positive for Ki67 is the ratio of the number of Ki67 positive nuclei over the total number of nuclei.

### ELISA

Protein levels of IL-6, IL-8, KIM-1 (HAVCR1), total MMP1, and total MMP7 were quantified from device effluents using the DuoSet^®^ line of ELISAs from R&D Systems according to the manufacturer’s instructions. For quantification of MMP7, all samples were diluted 1:12 in reagent diluent (R&D systems, DY995). 50 pg/mL MMP1 was detected in 2% human serum and this value was subtracted from all serum-treated samples.

### RNA-seq data generation and analysis

For MPS cultures, the cells were harvested from devices by injecting 100 μL of detergent (Abcam, part 8206000) into the injection port using a 1mL slip-tip syringe (BD, 309659) equipped with a 22-gauge needle (BD, 305142). Cell lysate was collected into 900 μL Trizol and frozen at −80 degrees Celsius until extraction. RNA was isolated using a RNeasy Micro Kit (Qiagen, 74004) and the RNA library was prepared with the SMARTer Stranded Total RNA Sample Prep Kit - Low Input Mammalian (Takara catalog number 634861) and sequenced as described previously (*29*).

RNA-seq data for primary cultures of human tubules grown in 10% FBS were previously generated (*85*) and downloaded from the Gene Expression Omnibus (GSE115961). For intact human kidney cortex and medulla, RNA extraction and RNA-seq was performed by GeneWiz (South Plainfield, New Jersey). For both sets of RNA-seq data, the paired end fastq files were trimmed and then aligned to human reference genome sequence hg38 using STAR 2.7.5b (doi:10.1093/bioinformatics/bts635) with parameters --outFilterIntronMotifs RemoveNoncanonical --outFilterMismatchNoverReadLmax 0.04 using annotation file Homo_sapiens.GRCh38.100.gtf downloaded from Ensembl database. The number of reads aligned to each exon (feature) was counted using *featureCounts* from subread package (doi:10.1093/bioinformatics/btt656). Multimapped reads were excluded. Differential expression of genes and read count normalization was performed using DESeq2 (doi:10.1186/s13059-014-0550-8) with the design formula “∼ condition”. Identification of pathway perturbation and key upstream regulators was performed using Advaita iPathwayGuide (https://advaitabio.com/ipathwayguide/). Gene ontology and known transcription factor target enrichments using ENRICHR were both performed using the BioJupies web tool (https://doi.org/10.1016/j.cels.2018.10.007).

Slight variations of the above protocol and analysis were used for the albumin treated MPS. Cells were isolated by injecting 100 μL buffer RLT (Qiagen, 79216) and the lysate was collected at the outflow port into an additional 900 μL of buffer RLT then frozen at −80°C until total RNA extraction using a Qiagen RNeasy Micro Kit. Total RNA was normalized to 2ng in a total volume of 9μl and then transcribed to cDNA in a dedicated PCR clean workstation using the SMART-Seq v4 Ultra Low Input RNA Kit (Takara Bio, Mountain View, CA). Sequencing libraries were constructed from cDNA using the SMARTer ThruPLEX DNA-Seq kit (Takara Bio). Final libraries were quantified, and library insert size distribution was checked using the Bioanalyzer (Agilent Technologies, Santa Clara, CA). Base calls generated in real-time on the NovaSeq 6000 instrument (RTA 3.1.5); demultiplexed, unaligned BAM files produced by Picard ExtractIlluminaBarcodes and IlluminaBasecallsToSam were converted to FASTQ format using SamTools bam2fq (v1.4).

Resulting sequences were aligned to the reference genome GENCODE human release 30 using STAR (v2.6.1d). Aligned data were read into R (version 3.6.1) and summarized as counts per gene using the Bioconductor GenomicAlignments package (*87*). Before fitting any models, we first excluded any genes that were expressed at consistently low levels across all samples. Prior to filtering, we had 58,870 genes and after filtering we had data for 15,336 genes. We then performed a trimmed mean of M-values (TMM) normalization (*88*). We used the voom method from the Bioconductor limma package, which estimates the mean-variance relationship of the log2-counts per million (logCPM), and generates a precision weight for each observation and enters these into the limma analysis pipeline. We used the linear mixed model approach, fitting the treatment as the fixed effect and the donor as the random effect by estimating the within-donor correlation. We then fit a linear model with treatment and incorporating the within-donor correlation. Since not all donors received all the treatment at each condition, the mixed model approach gives more statistical power for the unbalanced design. Rather than using a post hoc fold change filtering criterion, we used TREAT approach (*89*), which incorporates the fold-change into the statistic, meaning that instead of testing for genes which have fold-changes different from zero, we tested whether the fold-change is greater than 1.1-fold in absolute value. We selected genes based on a threshold of 1.1-fold-change and a false discovery rate of 5%.

### Chromatin accessibility data

DNase-seq data for primary cultures of human tubules grown in 10% FBS were previously generated (*85*) and downloaded from the Gene Expression Omnibus (GSE115961). ATAC-seq was performed on snap-frozen samples of human kidney cortex by ActiveMotif (Carlsbad, California) and aligned to the GRCh38 (hg38) reference genome using BWA and default settings. Read depth was normalized by random downsampling to the level of the sample with lowest coverage (29,795,703 tags). Total peaks were called using MACS 2.1.0 at a cutoff of p-value 1e-7, without control file, and with the -nomodel option. Peak filtering was performed by removing false ChIP-Seq peaks as defined within the ENCODE blacklist. The resulting intervals were merged into a non-overlapping ATAC-seq masterlist. We then merged this with our previously reported masterlist (primary cultures of human kidney tubules, glomerular outgrowths and ENCODE tubule culture datasets RPTEC, HRE and HRCE) using a bedops −u command. Overlaps between lists of regulatory elements were determined using the bedops −e (element of) and bedmap set operations. To identify regulatory elements with significantly different accessibility between intact renal cortex and cultured tubules, we counted reads mapping to every master list element in each sample. We then utilized the DESeq2 software package in R to identify regulatory elements with significantly different accessibility between replicate cortex and tubule culture samples, analyzing each sample separately. Sites that met an adjusted p-value<0.001 were considered differentially accessible regulatory elements. The distance of regulatory elements to the nearest gene transcription start site (TSS) and the associated biologic ontologies were computed using the Stanford GREAT analysis tool. An adjusted p-value cutoff of <0.0001 was used for determination of differentially accessible regulatory elements within 2.5kb of the TSS of differentially expressed genes (also determined using an adjusted p-value<0.0001). Transcription factor motif enrichment analysis was performed using HOMER using the list of non-significantly changed regulatory elements as the background set and default settings. The 733-cell regulatory element index and UCSC custom track of transcription factor archetypes and DNaseI footprints were obtained from recent publications (*37, 38*).

## Supplementary Figures

**Supplementary Figure 1.**
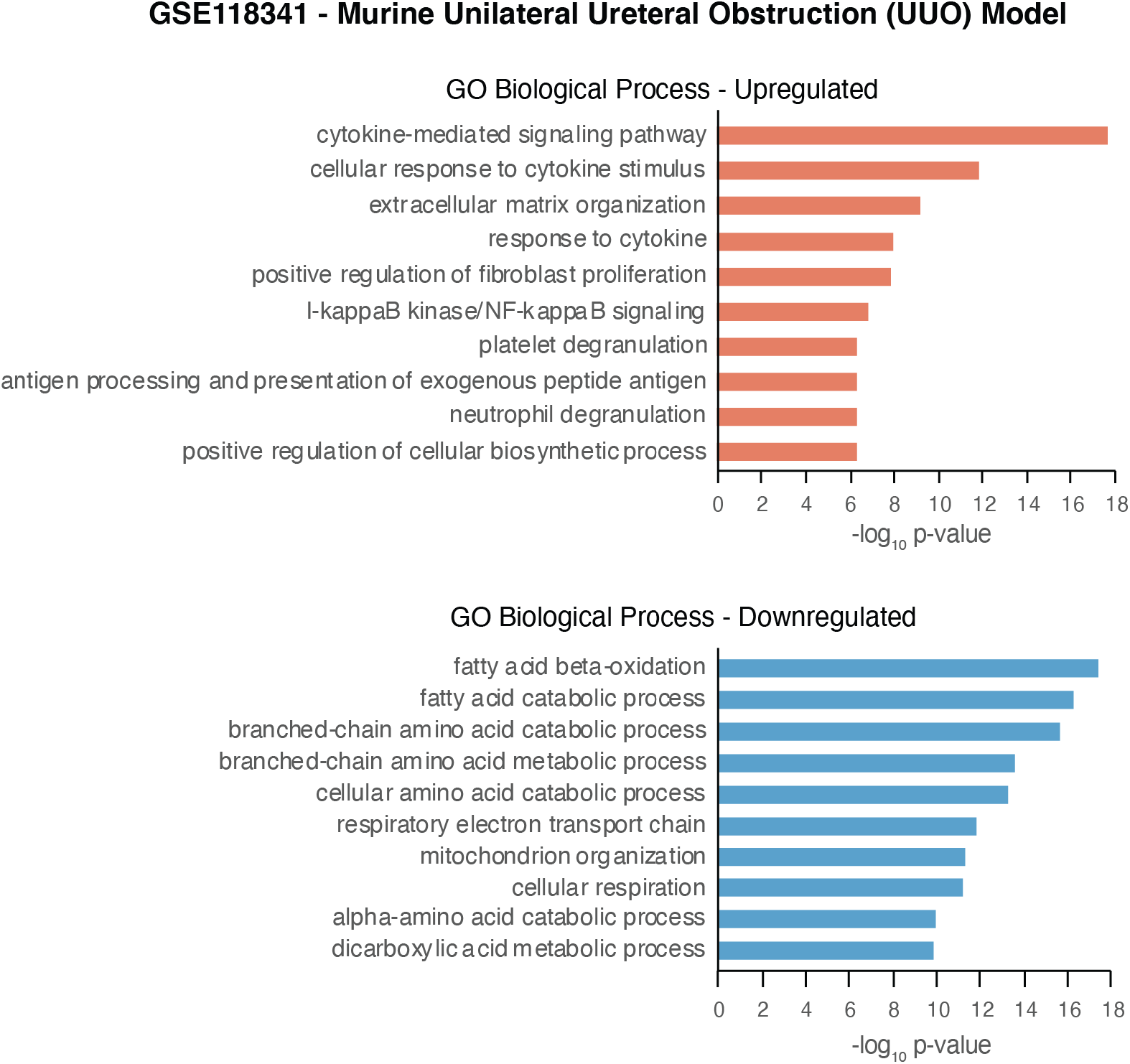
Gene ontology molecular function enrichments for differentially expressed genes in the murine unilateral ureteral obstruction model of tubular injury and fibrosis. From GSE118341. Compare to Figure 2B.

**Supplementary Figure 2.**
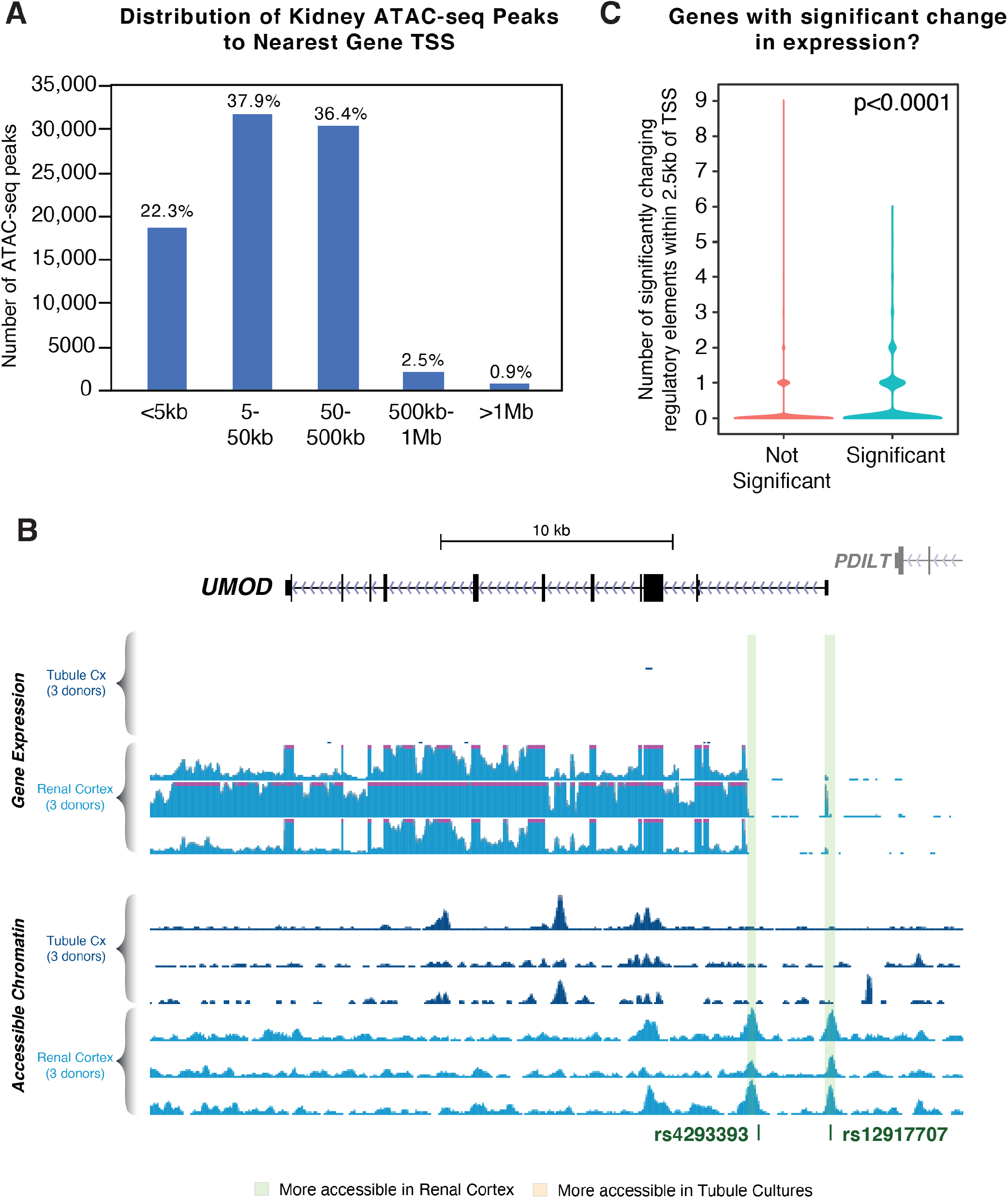
A) Distribution of the distance of intact kidney regulatory elements to the nearest gene transcription start site. B) Mapping of additional kidney disease GWAS loci to open chromatin regions in *UMOD* in adult kidney. Overview of the *UMOD* gene locus with gene expression (top) and accessible chromatin (bottom) tracks for cultured tubules and intact renal cortex (3 donors each). The *UMOD* gene is expressed at higher levels in intact renal cortex compared to tubules cultured in 10% FBS. Green vertical bars indicate two regulatory elements with greater accessibility in intact cortex which overlap kidney disease GWAS SNPs rs4293393 and rs12917707. C) Significantly changing genes are more likely to have a differentially accessible open chromatin region within 2.5kb of their TSS. p<0.0001 Chi-squared test with Yates correction.

**Supplementary Figure 3.**
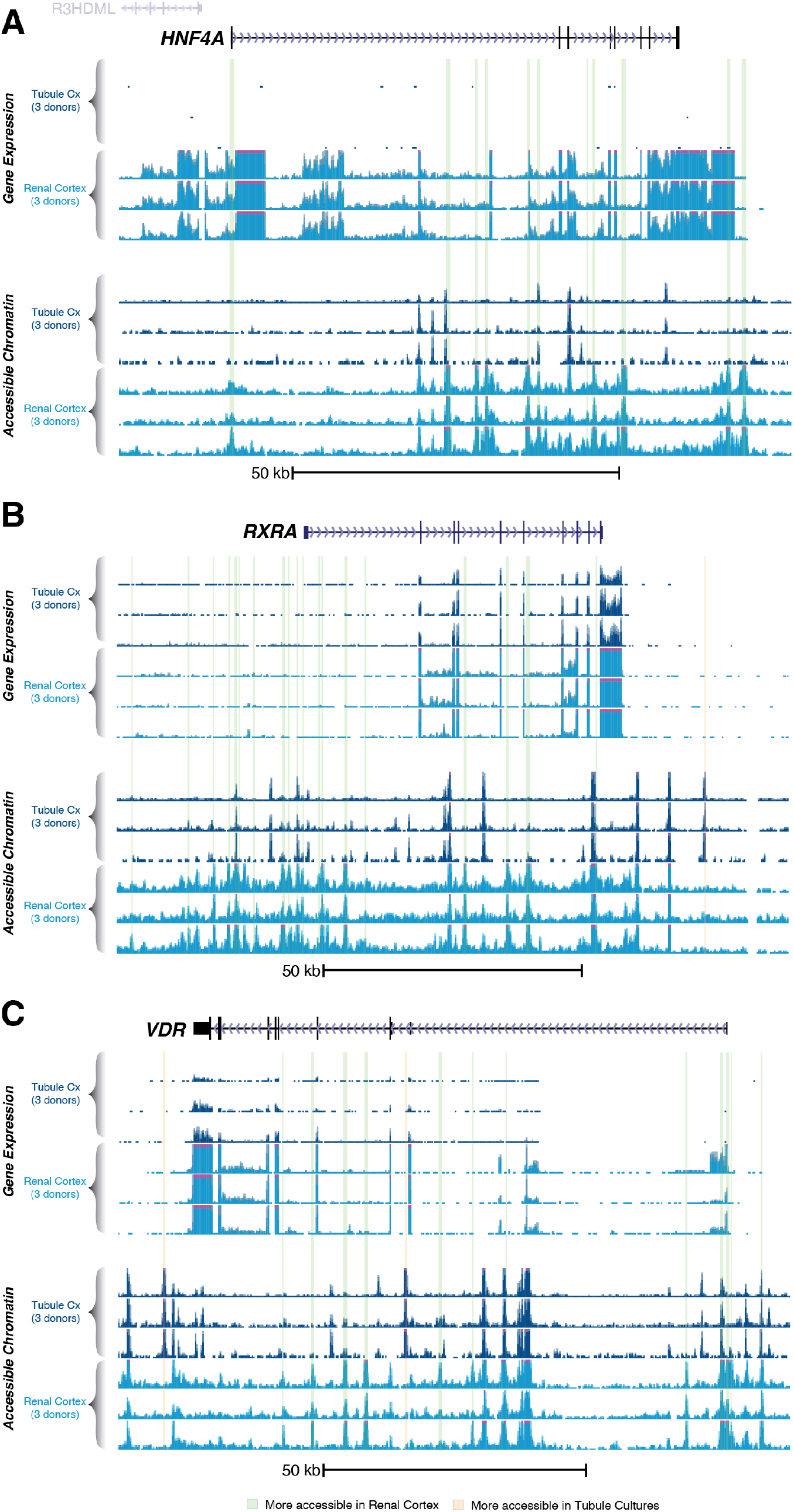
Regulatory landscape of key transcription factors/nuclear receptors. Overview of the A) *HNF4A* B) *RXRA* and C) *VDR* gene loci with gene expression (top) and accessible chromatin (bottom) tracks for cultured tubules and intact renal cortex (3 donors each). All 3 genes are expressed at higher levels in intact renal cortex compared to tubules cultured in 10% FBS. Green vertical bars indicate regulatory elements with greater accessibility in intact cortex.

**Supplementary Figure 4.**
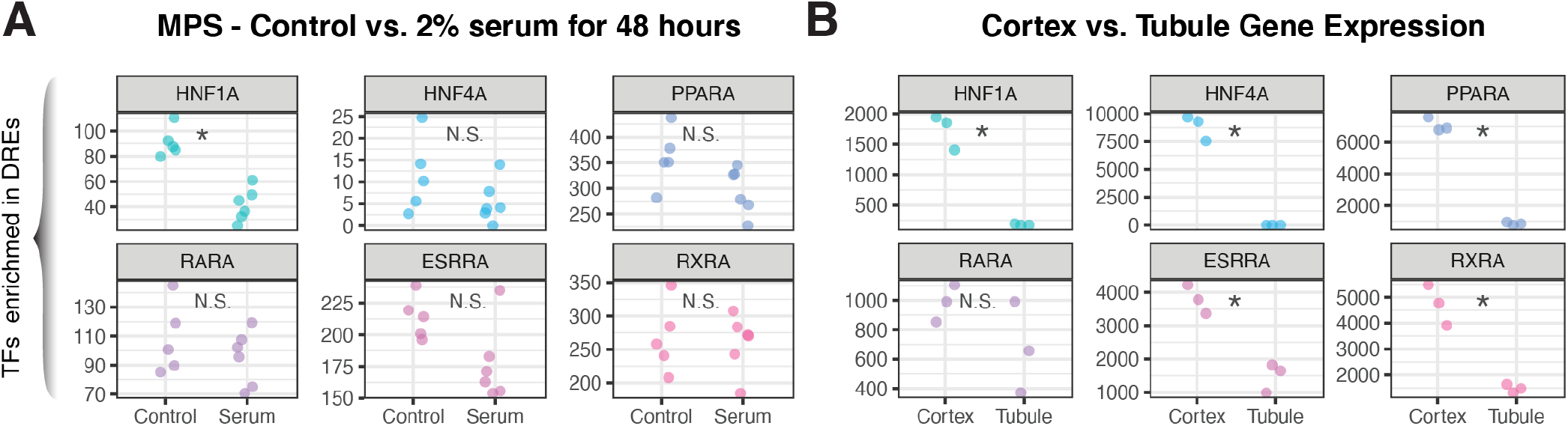
Expression of transcription factors associated with significant motif enrichments in intact renal cortex. Expression of selected transcription factors is shown in A) serum-treated tubule MPS and B) tubules cultured in 10% FBS.

**Supplementary Figure 5.**
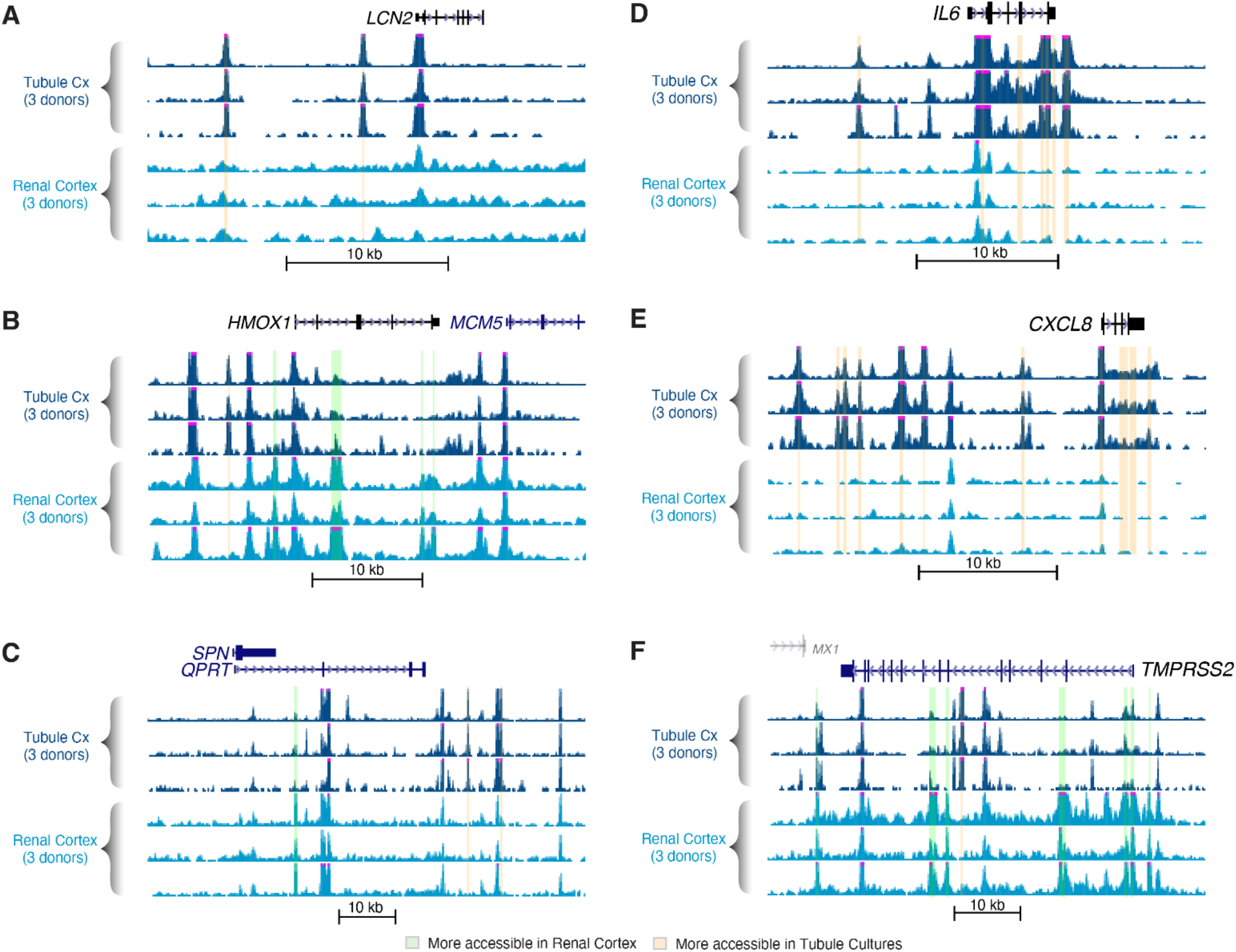
Regulatory landscape of kidney injury biomarker and COVID-19 associated genes. Overview of the indicated loci with gene expression (top) and accessible chromatin (bottom) tracks for cultured tubules and intact renal cortex (3 donors each). For kidney injury biomarkers, *LCN2* and *HMOX1* are expressed at higher levels in tubules cultured in 10% FBS compared to intact renal cortex whereas *QPRT* shows the opposite pattern of expression. For COVID-19 associated genes, *IL6* and *CXCL8* are expressed at higher levels in tubules cultured in 10% FBS compared to intact renal cortex whereas *TMPRSS2* shows the opposite pattern of expression. Green vertical bars indicate regulatory elements with greater accessibility in intact cortex, while orange vertical bars indicate regulatory elements with greater accessibility in cultured tubules.

**Supplementary Figure 6.**
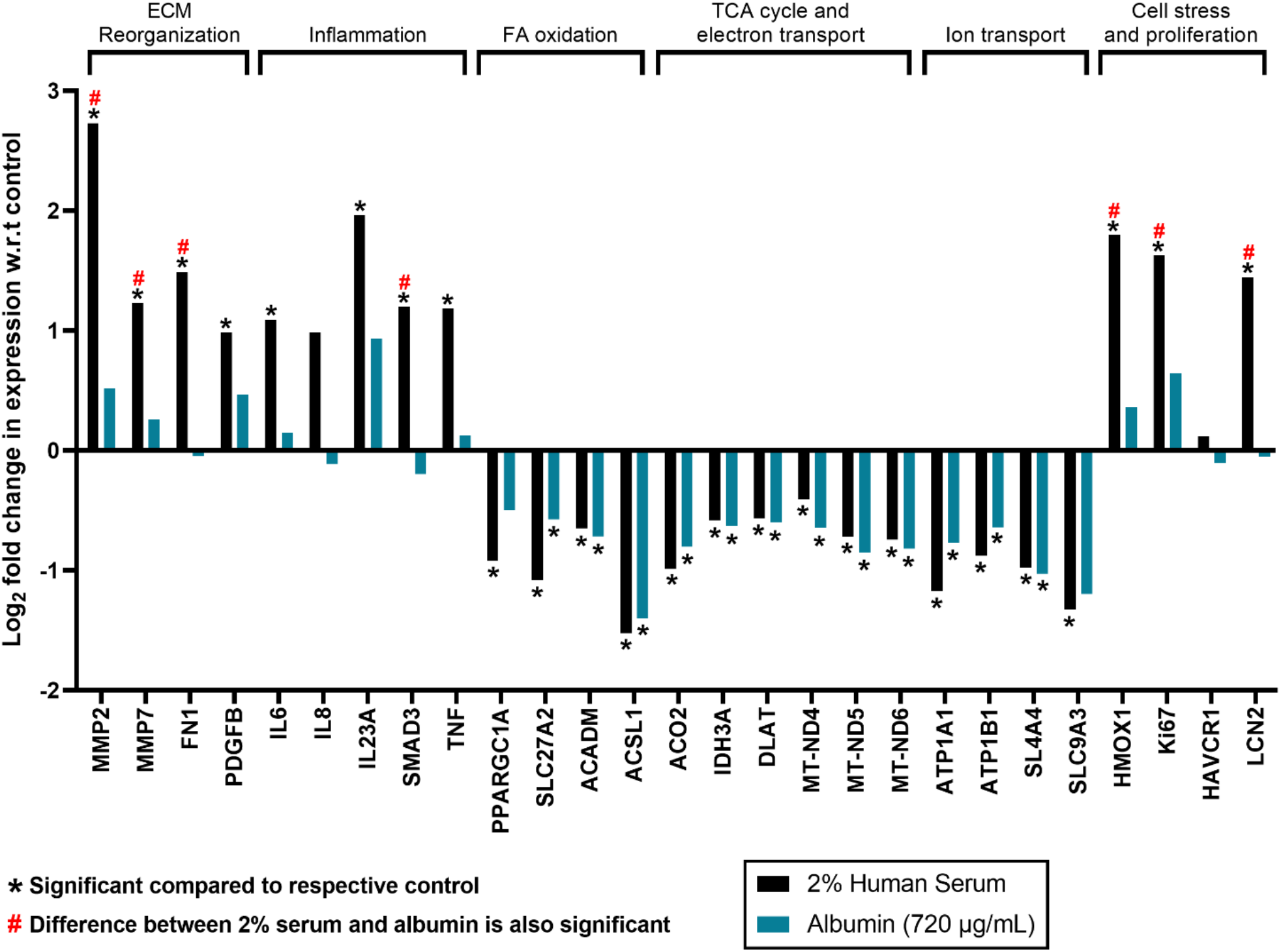
Albumin induces metabolic reprogramming, but not an inflammatory transcriptional signature in PTEC-MPS. MPS cultured PTECs were treated for 7 days with either 2% human serum or an equivalent concentration (720μg/ml) of albumin then harvested for RNA-seq. A selected panel of genes and the log_2_ fold change in transcript expression relative to control are shown. * indicates the expression of the indicated gene (serum or albumin-treated) is significantly different from the control sample. # indicates the difference between serum and albumin is statistically significant.

**Supplementary Table 1.**
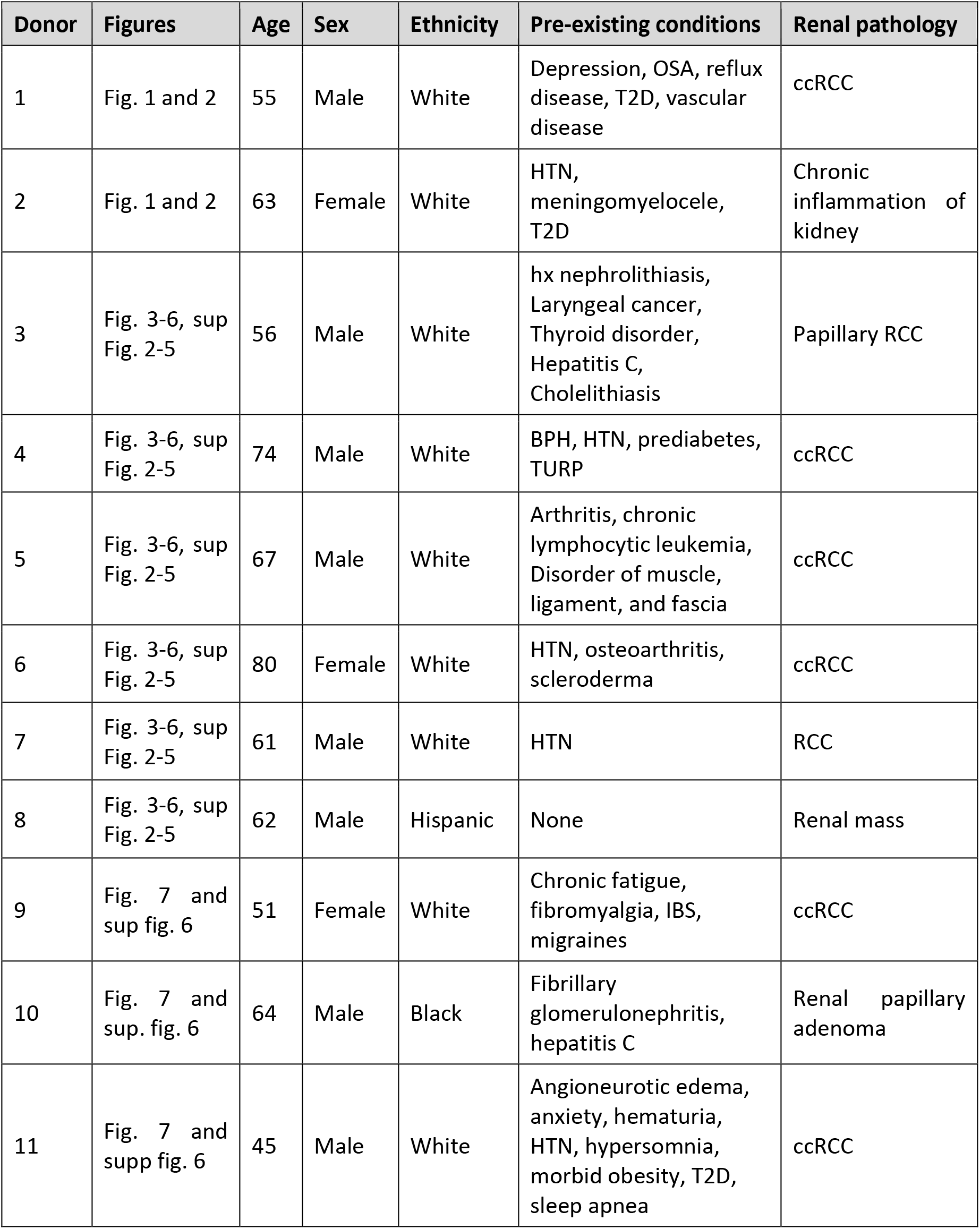
Demographics of donors from which cells or tissue were isolated. Abbreviations: BPH, benign prostatic hyperplasia; ccRCC, clear cell renal cell carcinoma; IBS, irritable bowel syndrome; hx, history of; HTN, hypertension OSA, obstructive sleep apnea; T2D, Type 2 diabetes; TURP, transurethral resection of the prostate.

## References

1. S. K. Mujais, K. Story, J. Brouillette, T. Takano, S. Soroka, C. Franek, D. Mendelssohn, F. O. Finkelstein, Health-related quality of life in CKD Patients: correlates and evolution over time. Clin J Am Soc Nephrol 4, 1293–1301 (2009).

2. M. Tonelli, N. Wiebe, B. Culleton, A. House, C. Rabbat, M. Fok, F. McAlister, A. X. Garg, Chronic kidney disease and mortality risk: a systematic review. J Am Soc Nephrol 17, 2034–2047 (2006).

3. U. S. R. D. System, 2019 USRDS annual data report: epidemiology of kidney disease in the United States., (2019).

4. L. S. Chawla, P. L. Kimmel, Acute kidney injury and chronic kidney disease: an integrated clinical syndrome. Kidney Int 82, 516–524 (2012).

5. L. A. Inker, A. S. Levey, K. Pandya, N. Stoycheff, A. Okparavero, T. Greene, C. Chronic Kidney Disease Epidemiology, Early change in proteinuria as a surrogate end point for kidney disease progression: an individual patient meta-analysis. Am J Kidney Dis 64, 74–85 (2014).

6. A. Thompson, K. Carroll, A. I. L, J. Floege, V. Perkovic, S. Boyer-Suavet, W. M. R, I. S. J, J. Barratt, D. C. Cattran, S. G. B, A. Kausz, W. M. A, H. N. Reich, H. R. B, M. West, P. H. Nachman, Proteinuria Reduction as a Surrogate End Point in Trials of IgA Nephropathy. Clin J Am Soc Nephrol 14, 469–481 (2019).

7. J. P. Troost, H. Trachtman, C. Spino, F. J. Kaskel, A. Friedman, M. M. Moxey-Mims, R. N. Fine, J. J. Gassman, J. B. Kopp, L. Walsh, R. Wang, D. S. Gipson, Proteinuria Reduction and Kidney Survival in Focal Segmental Glomerulosclerosis. Am J Kidney Dis, (2020).

8. M. E. Thomas, N. J. Brunskill, K. P. Harris, E. Bailey, J. H. Pringle, P. N. Furness, J. Walls, Proteinuria induces tubular cell turnover: A potential mechanism for tubular atrophy. Kidney Int 55, 890–898 (1999).

9. G. S. Hill, M. Delahousse, D. Nochy, C. Mandet, J. Bariety, Proteinuria and tubulointerstitial lesions in lupus nephritis. Kidney Int 60, 1893–1903 (2001).

10. A. Benigni, E. Gagliardini, A. Remuzzi, D. Corna, G. Remuzzi, Angiotensin-converting enzyme inhibition prevents glomerular-tubule disconnection and atrophy in passive Heymann nephritis, an effect not observed with a calcium antagonist. Am J Pathol 159, 1743–1750 (2001).

11. Y. Wang, G. K. Rangan, Y. C. Tay, Y. Wang, D. C. Harris, Induction of monocyte chemoattractant protein-1 by albumin is mediated by nuclear factor kappaB in proximal tubule cells. J Am Soc Nephrol 10, 1204–1213 (1999).

12. R. J. Baines, N. J. Brunskill, Tubular toxicity of proteinuria. Nat Rev Nephrol 7, 177–180 (2011).

13. B. A. Yard, E. Chorianopoulos, D. Herr, F. J. van der Woude, Regulation of endothelin-1 and transforming growth factor-beta1 production in cultured proximal tubular cells by albumin and heparan sulphate glycosaminoglycans. Nephrol Dial Transplant 16, 1769–1775 (2001).

14. S. Tang, J. C. Leung, K. Abe, K. W. Chan, L. Y. Chan, T. M. Chan, K. N. Lai, Albumin stimulates interleukin-8 expression in proximal tubular epithelial cells in vitro and in vivo. J Clin Invest 111, 515–527 (2003).

15. S. Yung, R. C. Tsang, Y. Sun, J. K. Leung, T. M. Chan, Effect of human anti-DNA antibodies on proximal renal tubular epithelial cell cytokine expression: implications on tubulointerstitial inflammation in lupus nephritis. J Am Soc Nephrol 16, 3281–3294 (2005).

16. S. A. Mezzano, M. Barria, M. A. Droguett, M. E. Burgos, L. G. Ardiles, C. Flores, J. Egido, Tubular NF-kappaB and AP-1 activation in human proteinuric renal disease. Kidney Int 60, 1366–1377 (2001).

17. Y. Motoyoshi, T. Matsusaka, A. Saito, I. Pastan, T. E. Willnow, S. Mizutani, I. Ichikawa, Megalin contributes to the early injury of proximal tubule cells during nonselective proteinuria. Kidney Int 74, 1262–1269 (2008).

18. D. Liu, Y. Wen, T. T. Tang, L. L. Lv, R. N. Tang, H. Liu, K. L. Ma, S. D. Crowley, B. C. Liu, Megalin/Cubulin-Lysosome-mediated Albumin Reabsorption Is Involved in the Tubular Cell Activation of NLRP3 Inflammasome and Tubulointerstitial Inflammation. J Biol Chem 290, 18018–18028 (2015).

19. M. Abbate, C. Zoja, G. Remuzzi, How does proteinuria cause progressive renal damage? J Am Soc Nephrol 17, 2974–2984 (2006).

20. N. P. Curthoys, O. W. Moe, Proximal tubule function and response to acidosis. Clin J Am Soc Nephrol 9, 1627–1638 (2014).

21. B. D. Humphreys, S. Czerniak, D. P. DiRocco, W. Hasnain, R. Cheema, J. V. Bonventre, Repair of injured proximal tubule does not involve specialized progenitors. Proc Natl Acad Sci U S A 108, 9226–9231 (2011).

22. T. Kusaba, M. Lalli, R. Kramann, A. Kobayashi, B. D. Humphreys, Differentiated kidney epithelial cells repair injured proximal tubule. Proc Natl Acad Sci U S A 111, 1527–1532 (2014).

23. M. Chang-Panesso, F. F. Kadyrov, M. Lalli, H. Wu, S. Ikeda, E. Kefaloyianni, M. M. Abdelmageed, A. Herrlich, A. Kobayashi, B. D. Humphreys, FOXM1 drives proximal tubule proliferation during repair from acute ischemic kidney injury. J Clin Invest 129, 5501–5517 (2019).

24. H. M. Kang, S. H. Ahn, P. Choi, Y. A. Ko, S. H. Han, F. Chinga, A. S. Park, J. Tao, K. Sharma, J. Pullman, E. P. Bottinger, I. J. Goldberg, K. Susztak, Defective fatty acid oxidation in renal tubular epithelial cells has a key role in kidney fibrosis development. Nat Med 21, 37–46 (2015).

25. L. Yang, T. Y. Besschetnova, C. R. Brooks, J. V. Shah, J. V. Bonventre, Epithelial cell cycle arrest in G2/M mediates kidney fibrosis after injury. Nat Med 16, 535–543, 531p following 143 (2010).

26. H. Wu, C. F. Lai, M. Chang-Panesso, B. D. Humphreys, Proximal Tubule Translational Profiling during Kidney Fibrosis Reveals Proinflammatory and Long Noncoding RNA Expression Patterns with Sexual Dimorphism. J Am Soc Nephrol 31, 23–38 (2020).

27. I. Grgic, G. Campanholle, V. Bijol, C. Wang, V. S. Sabbisetti, T. Ichimura, B. D. Humphreys, J. V. Bonventre, Targeted proximal tubule injury triggers interstitial fibrosis and glomerulosclerosis. Kidney Int 82, 172–183 (2012).

28. E. J. Weber, A. Chapron, B. D. Chapron, J. L. Voellinger, K. A. Lidberg, C. K. Yeung, Z. Wang, Y. Yamaura, D. W. Hailey, T. Neumann, D. D. Shen, K. E. Thummel, K. A. Muczynski, J. Himmelfarb, E. J. Kelly, Development of a microphysiological model of human kidney proximal tubule function. Kidney Int 90, 627–637 (2016).

29. E. J. Weber, K. A. Lidberg, L. Wang, T. K. Bammler, J. W. MacDonald, M. J. Li, M. Redhair, W. M. Atkins, C. Tran, K. M. Hines, J. Herron, L. Xu, M. B. Monteiro, S. Ramm, V. Vaidya, M. Vaara, T. Vaara, J. Himmelfarb, E. J. Kelly, Human kidney on a chip assessment of polymyxin antibiotic nephrotoxicity. JCI Insight 3, (2018).

30. B. D. Chapron, A. Chapron, B. Phillips, M. C. Okoli, D. D. Shen, E. J. Kelly, J. Himmelfarb, K. E. Thummel, Reevaluating the role of megalin in renal vitamin D homeostasis using a human cell-derived microphysiological system. ALTEX 35, 504–515 (2018).

31. T. Imaoka, J. Yang, L. Wang, M. G. McDonald, Z. Afsharinejad, T. K. Bammler, K. Van Ness, C. K. Yeung, A. E. Rettie, J. Himmelfarb, E. J. Kelly, Microphysiological system modeling of ochratoxin A-associated nephrotoxicity. Toxicology 444, 152582 (2020).

32. S. Y. Chang, E. J. Weber, V. S. Sidorenko, A. Chapron, C. K. Yeung, C. Gao, Q. Mao, D. Shen, J. Wang, T. A. Rosenquist, K. G. Dickman, T. Neumann, A. P. Grollman, E. J. Kelly, J. Himmelfarb, D. L. Eaton, Human liver-kidney model elucidates the mechanisms of aristolochic acid nephrotoxicity. JCI Insight 2, (2017).

33. K. A. Homan, D. B. Kolesky, M. A. Skylar-Scott, J. Herrmann, H. Obuobi, A. Moisan, J. A. Lewis, Bioprinting of 3D Convoluted Renal Proximal Tubules on Perfusable Chips. Sci Rep 6, 34845 (2016).

34. N. Y. C. Lin, K. A. Homan, S. S. Robinson, D. B. Kolesky, N. Duarte, A. Moisan, J. A. Lewis, Renal reabsorption in 3D vascularized proximal tubule models. Proc Natl Acad Sci U S A 116, 5399–5404 (2019).

35. K. J. Jang, A. P. Mehr, G. A. Hamilton, L. A. McPartlin, S. Chung, K. Y. Suh, D. E. Ingber, Human kidney proximal tubule-on-a-chip for drug transport and nephrotoxicity assessment. Integr Biol (Camb) 5, 1119–1129 (2013).

36. M. Pavkovic, L. Pantano, C. V. Gerlach, S. Brutus, S. A. Boswell, R. A. Everley, J. V. Shah, S. H. Sui, V. S. Vaidya, Multi omics analysis of fibrotic kidneys in two mouse models. Sci Data 6, 92 (2019).

37. W. Meuleman, A. Muratov, E. Rynes, J. Halow, K. Lee, D. Bates, M. Diegel, D. Dunn, F. Neri, A. Teodosiadis, A. Reynolds, E. Haugen, J. Nelson, A. Johnson, M. Frerker, M. Buckley, R. Sandstrom, J. Vierstra, R. Kaul, J. Stamatoyannopoulos, Index and biological spectrum of human DNase I hypersensitive sites. Nature 584, 244–251 (2020).

38. J. Vierstra, J. Lazar, R. Sandstrom, J. Halow, K. Lee, D. Bates, M. Diegel, D. Dunn, F. Neri, E. Haugen, E. Rynes, A. Reynolds, J. Nelson, A. Johnson, M. Frerker, M. Buckley, R. Kaul, W. Meuleman, J. A. Stamatoyannopoulos, Global reference mapping of human transcription factor footprints. Nature 583, 729–736 (2020).

39. P. Puigserver, B. M. Spiegelman, Peroxisome proliferator-activated receptor-gamma coactivator 1 alpha (PGC-1 alpha): transcriptional coactivator and metabolic regulator. Endocr Rev 24, 78–90 (2003).

40. M. Tran, D. Tam, A. Bardia, M. Bhasin, G. C. Rowe, A. Kher, Z. K. Zsengeller, M. R. Akhavan-Sharif, E. V. Khankin, M. Saintgeniez, S. David, D. Burstein, S. A. Karumanchi, I. E. Stillman, Z. Arany, S. M. Parikh, PGC-1alpha promotes recovery after acute kidney injury during systemic inflammation in mice. J Clin Invest 121, 4003–4014 (2011).

41. M. Fontecha-Barriuso, D. Martin-Sanchez, J. M. Martinez-Moreno, S. Carrasco, O. Ruiz-Andres, M. Monsalve, C. Sanchez-Ramos, M. J. Gomez, M. Ruiz-Ortega, M. D. Sanchez-Nino, P. Cannata-Ortiz, R. Cabello, C. Gonzalez-Enguita, A. Ortiz, A. B. Sanz, PGC-1alpha deficiency causes spontaneous kidney inflammation and increases the severity of nephrotoxic AKI. J Pathol 249, 65–78 (2019).

42. M. T. Tran, Z. K. Zsengeller, A. H. Berg, E. V. Khankin, M. K. Bhasin, W. Kim, C. B. Clish, I. E. Stillman, S. A. Karumanchi, E. P. Rhee, S. M. Parikh, PGC1alpha drives NAD biosynthesis linking oxidative metabolism to renal protection. Nature 531, 528–532 (2016).

43. A. Poyan Mehr, M. T. Tran, K. M. Ralto, D. E. Leaf, V. Washco, J. Messmer, A. Lerner, A. Kher, S. H. Kim, C. C. Khoury, S. J. Herzig, M. E. Trovato, N. Simon-Tillaux, M. R. Lynch, R. I. Thadhani, C. B. Clish, K. R. Khabbaz, E. P. Rhee, S. S. Waikar, A. H. Berg, S. M. Parikh, De novo NAD(+) biosynthetic impairment in acute kidney injury in humans. Nat Med 24, 1351–1359 (2018).

44. A. Tin, J. Marten, V. L. Halperin Kuhns, Y. Li, M. Wuttke, H. Kirsten, K. B. Sieber, C. Qiu, M. Gorski, Z. Yu, A. Giri, G. Sveinbjornsson, M. Li, A. Y. Chu, A. Hoppmann, L. J. O’Connor, B. Prins, T. Nutile, D. Noce, M. Akiyama, M. Cocca, S. Ghasemi, P. J. van der Most, K. Horn, Y. Xu, C. Fuchsberger, S. Sedaghat, S. Afaq, N. Amin, J. Arnlov, S. J. L. Bakker, N. Bansal, D. Baptista, S. Bergmann, M. L. Biggs, G. Biino, E. Boerwinkle, E. P. Bottinger, T. S. Boutin, M. Brumat, R. Burkhardt, E. Campana, A. Campbell, H. Campbell, R. J. Carroll, E. Catamo, J. C. Chambers, M. Ciullo, M. P. Concas, J. Coresh, T. Corre, D. Cusi, S. C. Felicita, M. H. de Borst, A. De Grandi, R. de Mutsert, A. P. J. de Vries, G. Delgado, A. Demirkan, O. Devuyst, K. Dittrich, K. U. Eckardt, G. Ehret, K. Endlich, M. K. Evans, R. T. Gansevoort, P. Gasparini, V. Giedraitis, C. Gieger, G. Girotto, M. Gogele, S. D. Gordon, D. F. Gudbjartsson, V. Gudnason, S. German Chronic Kidney Disease, T. Haller, P. Hamet, T. B. Harris, C. Hayward, A. A. Hicks, E. Hofer, H. Holm, W. Huang, N. Hutri-Kahonen, S. J. Hwang, M. A. Ikram, R. M. Lewis, E. Ingelsson, J. Jakobsdottir, I. Jonsdottir, H. Jonsson, P. K. Joshi, N. S. Josyula, B. Jung, M. Kahonen, Y. Kamatani, M. Kanai, S. M. Kerr, W. Kiess, M. E. Kleber, W. Koenig, J. S. Kooner, A. Korner, P. Kovacs, B. K. Kramer, F. Kronenberg, M. Kubo, B. Kuhnel, M. La Bianca, L. A. Lange, B. Lehne, T. Lehtimaki, S. Lifelines Cohort, J. Liu, M. Loeffler, R. J. F. Loos, L. P. Lyytikainen, R. Magi, A. Mahajan, N. G. Martin, W. Marz, D. Mascalzoni, K. Matsuda, C. Meisinger, T. Meitinger, A. Metspalu, Y. Milaneschi, V. A. M. V. Program, C. J. O’Donnell, O. D. Wilson, J. M. Gaziano, P. P. Mishra, K. L. Mohlke, N. Mononen, G. W. Montgomery, D. O. Mook-Kanamori, M. Muller-Nurasyid, G. N. Nadkarni, M. A. Nalls, M. Nauck, K. Nikus, B. Ning, I. M. Nolte, R. Noordam, J. R. O’Connell, I. Olafsson, S. Padmanabhan, B. Penninx, T. Perls, A. Peters, M. Pirastu, N. Pirastu, G. Pistis, O. Polasek, B. Ponte, D. J. Porteous, T. Poulain, M. H. Preuss, T. J. Rabelink, L. M. Raffield, O. T. Raitakari, R. Rettig, M. Rheinberger, K. M. Rice, F. Rizzi, A. Robino, I. Rudan, A. Krajcoviechova, R. Cifkova, R. Rueedi, D. Ruggiero, K. A. Ryan, Y. Saba, E. Salvi, H. Schmidt, R. Schmidt, C. M. Shaffer, A. V. Smith, B. H. Smith, C. N. Spracklen, K. Strauch, M. Stumvoll, P. Sulem, S. M. Tajuddin, A. Teren, J. Thiery, C. H. L. Thio, U. Thorsteinsdottir, D. Toniolo, A. Tonjes, J. Tremblay, A. G. Uitterlinden, S. Vaccargiu, P. van der Harst, C. M. van Duijn, N. Verweij, U. Volker, P. Vollenweider, G. Waeber, M. Waldenberger, J. B. Whitfield, S. H. Wild, J. F. Wilson, Q. Yang, W. Zhang, A. B. Zonderman, M. Bochud, J. G. Wilson, S. A. Pendergrass, K. Ho, A. Parsa, P. P. Pramstaller, B. M. Psaty, C. A. Boger, H. Snieder, A. S. Butterworth, Y. Okada, T. L. Edwards, K. Stefansson, K. Susztak, M. Scholz, I. M. Heid, A. M. Hung, A. Teumer, C. Pattaro, O. M. Woodward, V. Vitart, A. Kottgen, Target genes, variants, tissues and transcriptional pathways influencing human serum urate levels. Nat Genet 51, 1459–1474 (2019).

45. S. Muthusamy, J. J. Jeong, M. Cheng, J. A. Bonzo, A. Kumar, F. J. Gonzalez, A. Borthakur, P. K. Dudeja, S. Saksena, J. Malakooti, Hepatocyte nuclear factor 4alpha regulates the expression of intestinal epithelial Na(+)/H(+) exchanger isoform 3. Am J Physiol Gastrointest Liver Physiol 314, G14–G21 (2018).

46. A. Joseph, R. A. Hess, D. J. Schaeffer, C. Ko, S. Hudgin-Spivey, P. Chambon, B. D. Shur, Absence of estrogen receptor alpha leads to physiological alterations in the mouse epididymis and consequent defects in sperm function. Biol Reprod 82, 948–957 (2010).

47. B. Li, R. A. Rietmeijer, S. G. Brohawn, Structural basis for pH gating of the two-pore domain K(+) channel TASK2. Nature 586, 457–462 (2020).

48. H. Barriere, R. Belfodil, I. Rubera, M. Tauc, F. Lesage, C. Poujeol, N. Guy, J. Barhanin, P. Poujeol, Role of TASK2 potassium channels regarding volume regulation in primary cultures of mouse proximal tubules. J Gen Physiol 122, 177–190 (2003).

49. R. Warth, H. Barriere, P. Meneton, M. Bloch, J. Thomas, M. Tauc, D. Heitzmann, E. Romeo, F. Verrey, R. Mengual, N. Guy, S. Bendahhou, F. Lesage, P. Poujeol, J. Barhanin, Proximal renal tubular acidosis in TASK2 K+ channel-deficient mice reveals a mechanism for stabilizing bicarbonate transport. Proc Natl Acad Sci U S A 101, 8215–8220 (2004).

50. D. Toncheva, M. Mihailova-Hristova, R. Vazharova, R. Staneva, S. Karachanak, P. Dimitrov, V. Simeonov, S. Ivanov, L. Balabanski, D. Serbezov, M. Malinov, V. Stefanovic, R. Cukuranovic, M. Polenakovic, L. Jankovic-Velickovic, V. Djordjevic, T. Jevtovic-Stoimenov, D. Plaseska-Karanfilska, A. Galabov, V. Djonov, I. Dimova, NGS nominated CELA1, HSPG2, and KCNK5 as candidate genes for predisposition to Balkan endemic nephropathy. Biomed Res Int 2014, 920723 (2014).

51. B. Zhang, G. Ramesh, C. C. Norbury, W. B. Reeves, Cisplatin-induced nephrotoxicity is mediated by tumor necrosis factor-alpha produced by renal parenchymal cells. Kidney Int 72, 37–44 (2007).

52. M. C. Wiggins, M. Bracher, A. Mall, R. Hickman, S. C. Robson, D. Kahn, Tumour necrosis factor levels during acute rejection and acute tubular necrosis in renal transplant recipients. Transpl Immunol 8, 211–215 (2000).

53. G. Ramesh, W. B. Reeves, TNF-alpha mediates chemokine and cytokine expression and renal injury in cisplatin nephrotoxicity. J Clin Invest 110, 835–842 (2002).

54. G. Guo, J. Morrissey, R. McCracken, T. Tolley, S. Klahr, Role of TNFR1 and TNFR2 receptors in tubulointerstitial fibrosis of obstructive nephropathy. Am J Physiol 277, F766–772 (1999).

55. O. Kwon, B. A. Molitoris, M. Pescovitz, K. J. Kelly, Urinary actin, interleukin-6, and interleukin-8 may predict sustained ARF after ischemic injury in renal allografts. Am J Kidney Dis 41, 1074–1087 (2003).

56. D. Zhou, Y. Tian, L. Sun, L. Zhou, L. Xiao, R. J. Tan, J. Tian, H. Fu, F. F. Hou, Y. Liu, Matrix Metalloproteinase-7 Is a Urinary Biomarker and Pathogenic Mediator of Kidney Fibrosis. J Am Soc Nephrol 28, 598–611 (2017).

57. X. Yang, C. Chen, S. Teng, X. Fu, Y. Zha, H. Liu, L. Wang, J. Tian, X. Zhang, Y. Liu, J. Nie, F. F. Hou, Urinary Matrix Metalloproteinase-7 Predicts Severe AKI and Poor Outcomes after Cardiac Surgery. J Am Soc Nephrol 28, 3373–3382 (2017).

58. H. Fu, D. Zhou, H. Zhu, J. Liao, L. Lin, X. Hong, F. F. Hou, Y. Liu, Matrix metalloproteinase-7 protects against acute kidney injury by priming renal tubules for survival and regeneration. Kidney Int 95, 1167–1180 (2019).

59. W. He, R. J. Tan, Y. Li, D. Wang, J. Nie, F. F. Hou, Y. Liu, Matrix metalloproteinase-7 as a surrogate marker predicts renal Wnt/beta-catenin activity in CKD. J Am Soc Nephrol 23, 294–304 (2012).

60. K. Surendran, T. C. Simon, H. Liapis, J. K. McGuire, Matrilysin (MMP-7) expression in renal tubular damage: association with Wnt4. Kidney Int 65, 2212–2222 (2004).

61. M. Afkarian, L. R. Zelnick, J. Ruzinski, B. Kestenbaum, J. Himmelfarb, I. H. de Boer, R. Mehrotra, Urine matrix metalloproteinase-7 and risk of kidney disease progression and mortality in type 2 diabetes. J Diabetes Complications 29, 1024–1031 (2015).

62. D. E. Oken, W. Flamenbaum, Micropuncture studies of proximal tubule albumin concentrations in normal and nephrotic rats. Journal of Clinical Investigation 50, 1498–1505 (1971).

63. H. Stolte, H. J. Schurek, J. M. Alt, Glomerular albumin filtration: a comparison of micropuncture studies in the isolated perfused rat kidney with in vivo experimental conditions. Kidney Int 16, 377–384 (1979).

64. J. Van Liew, W. Buentig, H. Stolte, J. Boylan, Protein excretion: micropuncture study of rat capsular and proximal tubule fluid. American Journal of Physiology-Legacy Content 219, 299–305 (1970).

65. A. Tojo, H. Endou, Intrarenal handling of proteins in rats using fractional micropuncture technique. Am J Physiol 263, F601–606 (1992).

66. L. M. Russo, R. M. Sandoval, S. B. Campos, B. A. Molitoris, W. D. Comper, D. Brown, Impaired Tubular Uptake Explains Albuminuria in Early Diabetic Nephropathy. 20, 489–494 (2009).

67. L. M. Russo, R. M. Sandoval, M. McKee, T. M. Osicka, A. B. Collins, D. Brown, B. A. Molitoris, W. D. Comper, The normal kidney filters nephrotic levels of albumin retrieved by proximal tubule cells: Retrieval is disrupted in nephrotic states. Kidney International 71, 504–513 (2007).

68. G. R. Joachim, J. S. Cameron, M. Schwartz, E. L. Becker, Selectivity of Protein Excretion in Patients with the Nephrotic Syndrome. J Clin Invest 43, 2332–2346 (1964).

69. S. Friedman, H. W. Jones, 3rd, H. V. Golbetz, J. A. Lee, H. L. Little, B. D. Myers, Mechanisms of proteinuria in diabetic nephropathy. II. A study of the size-selective glomerular filtration barrier. Diabetes 32 Suppl 2, 40–46 (1983).

70. B. J. Carrie, B. D. Myers, Proteinuria and functional characteristics of the glomerular barrier in diabetic nephropathy. Kidney Int 17, 669–676 (1980).

71. P. Lee, X. Wu, Review: modifications of human serum albumin and their binding effect. Curr Pharm Des 21, 1862–1865 (2015).

72. S. P. S. Yadav, R. M. Sandoval, J. Zhao, Y. Huang, E. Wang, S. Kumar, S. B. Campos-Bilderback, G. Rhodes, Y. Mechref, B. A. Molitoris, M. C. Wagner, Mechanism of how carbamylation reduces albumin binding to FcRn contributing to increased vascular clearance. Am J Physiol Renal Physiol 320, F114–F129 (2021).

73. A. Raghav, J. Ahmad, Glycated albumin in chronic kidney disease: Pathophysiologic connections. Diabetes Metab Syndr 12, 463–468 (2018).

74. A. R. Chade, M. L. Williams, J. E. Engel, E. Williams, G. L. Bidwell, 3rd, Molecular targeting of renal inflammation using drug delivery technology to inhibit NF-kappaB improves renal recovery in chronic kidney disease. Am J Physiol Renal Physiol 319, F139–F148 (2020).

75. G. K. Rangan, Y. Wang, Y. C. Tay, D. C. Harris, Inhibition of nuclear factor-kappaB activation reduces cortical tubulointerstitial injury in proteinuric rats. Kidney Int 56, 118–134 (1999).

76. G. Canaud, C. R. Brooks, S. Kishi, K. Taguchi, K. Nishimura, S. Magassa, A. Scott, L. L. Hsiao, T. Ichimura, F. Terzi, L. Yang, J. V. Bonventre, Cyclin G1 and TASCC regulate kidney epithelial cell G2-M arrest and fibrotic maladaptive repair. Sci Transl Med 11, (2019).

77. Z. Daniloski, T. X. Jordan, H.-H. Wessels, D. A. Hoagland, S. Kasela, M. Legut, S. Maniatis, E. P. Mimitou, L. Lu, E. Geller, O. Danziger, B. R. Rosenberg, H. Phatnani, P. Smibert, T. Lappalainen, B. R. Tenoever, N. E. Sanjana, Identification of Required Host Factors for SARS-CoV-2 Infection in Human Cells. Cell 184, 92–105.e116 (2021).

78. M. Singh, V. Bansal, C. Feschotte, A Single-Cell RNA Expression Map of Human Coronavirus Entry Factors. Cell Rep 32, 108175 (2020).

79. V. Monteil, H. Kwon, P. Prado, A. Hagelkruys, R. A. Wimmer, M. Stahl, A. Leopoldi, E. Garreta, C. Hurtado Del Pozo, F. Prosper, J. P. Romero, G. Wirnsberger, H. Zhang, A. S. Slutsky, R. Conder, N. Montserrat, A. Mirazimi, J. M. Penninger, Inhibition of SARS-CoV-2 Infections in Engineered Human Tissues Using Clinical-Grade Soluble Human ACE2. Cell 181, 905–913 e907 (2020).

80. L. Chan, K. Chaudhary, A. Saha, K. Chauhan, A. Vaid, S. Zhao, I. Paranjpe, S. Somani, F. Richter, R. Miotto, A. Lala, A. Kia, P. Timsina, L. Li, R. Freeman, R. Chen, J. Narula, A. C. Just, C. Horowitz, Z. Fayad, C. Cordon-Cardo, E. Schadt, M. A. Levin, D. L. Reich, V. Fuster, B. Murphy, J. C. He, A. W. Charney, E. P. Bottinger, B. S. Glicksberg, S. G. Coca, G. N. Nadkarni, C. I. C. Mount Sinai, AKI in Hospitalized Patients with COVID-19. J Am Soc Nephrol 32, 151–160 (2021).

81. D. Blanco-Melo, B. E. Nilsson-Payant, W. C. Liu, S. Uhl, D. Hoagland, R. Moller, T. X. Jordan, K. Oishi, M. Panis, D. Sachs, T. T. Wang, R. E. Schwartz, J. K. Lim, R. A. Albrecht, B. R. tenOever, Imbalanced Host Response to SARS-CoV-2 Drives Development of COVID-19. Cell 181, 1036–1045 e1039 (2020).

82. O. Takase, T. Marumo, N. Imai, J. Hirahashi, A. Takayanagi, K. Hishikawa, M. Hayashi, N. Shimizu, T. Fujita, T. Saruta, NF-kappaB-dependent increase in intrarenal angiotensin II induced by proteinuria. Kidney Int 68, 464–473 (2005).

83. B. C. Liu, J. Gao, Q. Li, L. M. Xu, Albumin caused the increasing production of angiotensin II due to the dysregulation of ACE/ACE2 expression in HK2 cells. Clin Chim Acta 403, 23–30 (2009).

84. M. A. Crackower, R. Sarao, G. Y. Oudit, C. Yagil, I. Kozieradzki, S. E. Scanga, A. J. Oliveira-dos-Santos, J. da Costa, L. Zhang, Y. Pei, J. Scholey, C. M. Ferrario, A. S. Manoukian, M. C. Chappell, P. H. Backx, Y. Yagil, J. M. Penninger, Angiotensin-converting enzyme 2 is an essential regulator of heart function. Nature 417, 822–828 (2002).

85. K. B. Sieber, A. Batorsky, K. Siebenthall, K. L. Hudkins, J. D. Vierstra, S. Sullivan, A. Sur, M. McNulty, R. Sandstrom, A. Reynolds, D. Bates, M. Diegel, D. Dunn, J. Nelson, M. Buckley, R. Kaul, M. G. Sampson, J. Himmelfarb, C. E. Alpers, D. Waterworth, S. Akilesh, Integrated Functional Genomic Analysis Enables Annotation of Kidney Genome-Wide Association Study Loci. J Am Soc Nephrol, (2019).

86. K. P. Van Ness, S. Y. Chang, E. J. Weber, D. Zumpano, D. L. Eaton, E. J. Kelly, Microphysiological Systems to Assess Nonclinical Toxicity. Curr Protoc Toxicol 73, 14 18 11–14 18 28 (2017).

87. M. Lawrence, W. Huber, H. Pages, P. Aboyoun, M. Carlson, R. Gentleman, M. T. Morgan, V. J. Carey, Software for computing and annotating genomic ranges. PLoS Comput Biol 9, e1003118 (2013).

88. M. D. Robinson, A. Oshlack, A scaling normalization method for differential expression analysis of RNA-seq data. Genome Biol 11, R25 (2010).

89. D. J. McCarthy, G. K. Smyth, Testing significance relative to a fold-change threshold is a TREAT. Bioinformatics 25, 765–771 (2009).

